# The human proteome co-regulation map reveals functional relationships between proteins

**DOI:** 10.1101/582247

**Authors:** Georg Kustatscher, Piotr Grabowski, Tina A. Schrader, Josiah B. Passmore, Michael Schrader, Juri Rappsilber

## Abstract

The annotation of protein function is a longstanding challenge of cell biology that suffers from the sheer magnitude of the task. Here we present ProteomeHD, which documents the response of 10,323 human proteins to 294 biological perturbations, measured by isotope-labelling mass spectrometry. Using this data matrix and robust machine learning we create a co-regulation map of the cell that reflects functional associations between human proteins. The map identifies a functional context for many uncharacterized proteins, including microproteins that are difficult to study with traditional methods. Co-regulation also captures relationships between proteins which do not physically interact or co-localize. For example, co-regulation of the peroxisomal membrane protein PEX11β with mitochondrial respiration factors led us to discover a novel organelle interface between peroxisomes and mitochondria in mammalian cells. The co-regulation map can be explored at www.proteomeHD.net.

---

Functional genomics approaches often use a “guilt-by-association” strategy to determine the biological function of genes and proteins on a system-wide scale. For example, high-throughput measurement of protein-protein interactions^1-5^ and subcellular localization^6-9^ has delivered invaluable insights into proteome organisation. A limitation of these techniques is that extensive biochemical procedures and cross-reacting antibodies may introduce artifacts. Moreover, not all proteins that function in the same biological process also interact physically or co-localize. Such functional relationships may be uncovered by assays with phenotypic readouts, including genetic interactions^10^ and metabolic profiles^11^, but these have yet to be applied on a genomic scale in humans. One of the oldest functional genomics methods is gene expression profiling^12^. Genes with correlated activity often participate in similar cellular functions, which can be exploited to infer the function of uncharacterized genes based on their coexpression with known genes^13-18^.

However, predicting gene function from coexpression alone often leads to inaccurate results^19,20^. One possible reason for this is that gene activity is generally measured at the mRNA level, neglecting the contribution of protein synthesis and degradation to gene expression control. The precise extent to which protein levels depend on mRNA abundances is still debated, and likely differs between genes and test systems^21-23^. However, some fundamental differences between mRNA and protein expression control have recently emerged. For example, many genes have coexpressed mRNAs due to their chromosomal proximity rather than any functional similarity^19,24-26^. Such non-functional mRNA coexpression results from stochastic transitions between active and inactive chromatin that affect wide genomic loci^24,25,27^, and transcriptional interference between closeby genes^25,28^. Importantly, coexpression of spatially close, but functionally unrelated genes is buffered at the protein level^19,25^. Protein abundances are also less affected than mRNA levels by genetic variation^29,30^, including variations in gene copy numbers^31-33^. Consequently, protein expression profiling outperforms mRNA expression profiling with regard to gene function prediction^19,20^. Protein-based profiling not only allows for a more accurate measurement of gene activity, but can determine additional aspects of a cell’s response to a perturbation, such as changes in protein localization and modification state. At the proteome level, expression profiling can therefore be extended to a more comprehensive protein covariation analysis.

Proof-of-principle studies by us and others have shown that protein covariation can be used to infer, for example, the composition of protein complexes and organelles^34-42^. However, these studies have focussed on relatively small sets of proteins or biological conditions, or used samples tailored to the analysis of specific cellular structures. In addition to the limited amount of data, coexpression analyses may be held back by the statistical tools used to pinpoint genes with similar activity. Coexpressed genes are commonly identified using Pearson’s correlation, which is restricted to linear correlations and susceptible to outliers. Machine-learning may offer an increase in sensitivity and specificity.

Despite the success of functional genomics, many human proteins remain uncharacterized, especially small proteins that are difficult to study by biochemical methods. The emergence of big proteomics data and new computational approaches could provide an opportunity to look at these proteins from a different angle. We wondered if protein covariation would assign functions to previously uncharacterized proteins or novel roles to characterized ones. The resulting resource is available at www.proteomeHD.net to generate hypotheses on the cellular functions of proteins of interest in a straightforward manner.

## RESULTS

### ProteomeHD is a data matrix for functional proteomics

To turn protein covariation analysis into a system-wide, generally applicable method, we created ProteomeHD. In contrast to previous drafts of the human proteome^8,9,22,43,44^, ProteomeHD does not catalogue the proteome of specific tissues or subcellular compartments. Instead, ProteomeHD catalogues the transitions between different proteome states, i.e. changes in protein abundance or localization resulting from cellular perturbations. HD, or high-definition, refers to two aspects of the dataset. First, all experiments are quantified using SILAC (stable isotope labelling by amino acids in cell culture)^45^. SILAC essentially eliminates sample processing artifacts and is especially accurate when quantifying small fold-changes. This is crucial to detect subtle, system-wide effects of a perturbation on the protein network. Second, HD refers to the number of observations (pixels) available for each protein. As more perturbations are analysed, regulatory patterns become more refined and can be detected more accurately.

To assemble ProteomeHD we processed the raw data from 5,288 individual mass-spectrometry runs into one coherent data matrix, which covers 10,323 proteins (from 9,987 genes) and 294 biological conditions (Supplementary Table 1). About 20% of the experiments were performed in our laboratory and the remaining data were collected from the Proteomics Identifications (PRIDE)^46^ repository (Fig. 1a). The data cover a wide array of quantitative proteomics experiments, such as perturbations with drugs and growth factors, genetic perturbations, cell differentiation studies and comparisons of cancer cell lines (Supplementary Table 2). All experiments are comparative studies using SILAC^45^, i.e. they do not report absolute protein concentrations but highly accurate fold-changes in response to perturbation. About 60% of the included experiments analysed whole-cell samples. The remaining measurements were performed on samples that had been fractionated after perturbation, e.g. to enrich for chromatin-based or secreted proteins. This allows for the detection of low-abundance proteins that may not be detected in whole-cell lysates.

**Figure 1.**
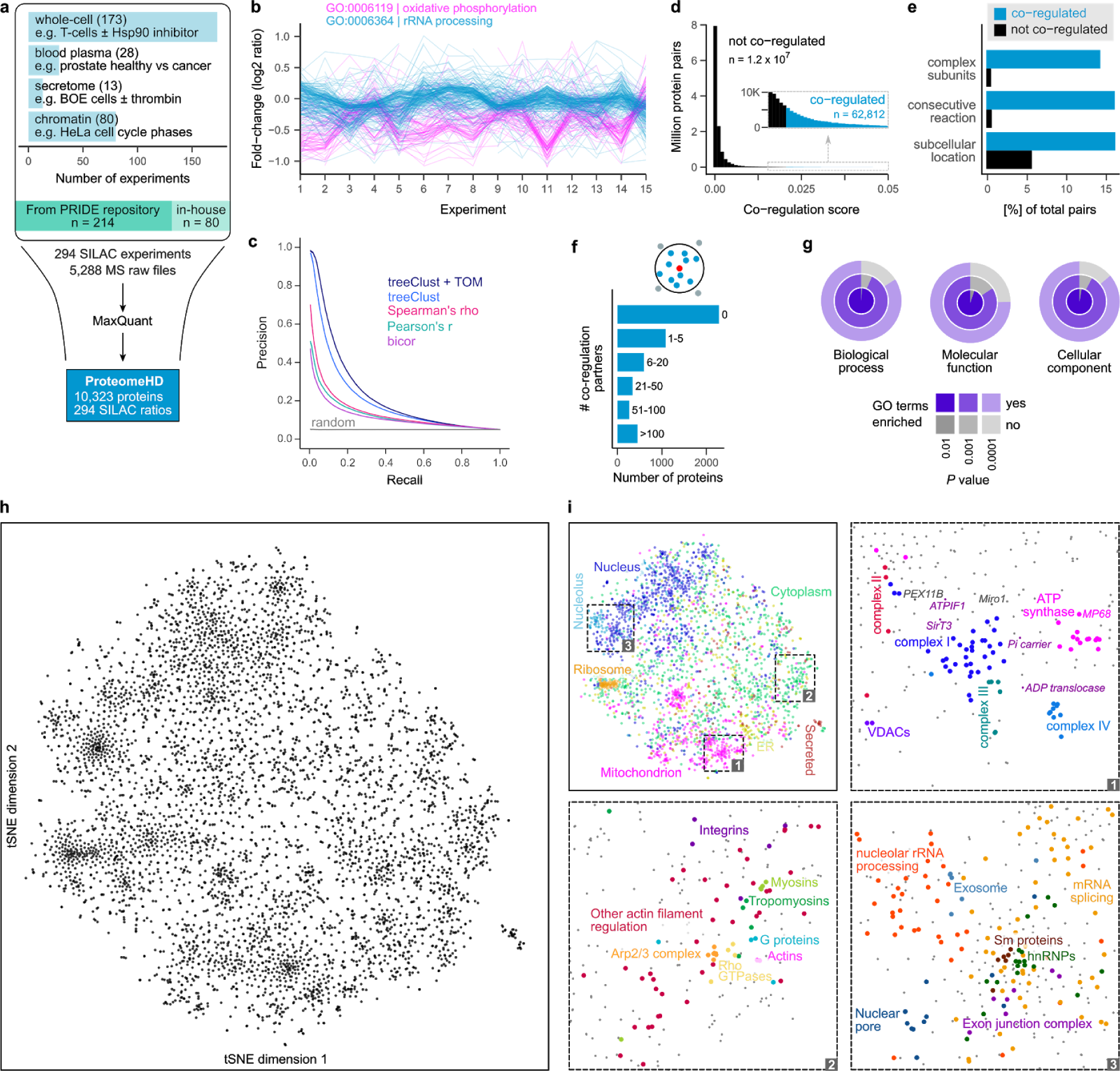
The co-regulation map shows functional associations between human proteins. (**a**) Assembly of ProteomeHD, which quantifies the protein response to 294 perturbations using SILAC^45^. Most measurements document protein abundance changes in whole-cell samples, but in some cases subcellular fractions were enriched to detect low-abundance proteins. Data were collected from PRIDE^46^ and produced in-house. (**b**) A random set of experiments from ProteomeHD, showing that groups of proteins with related functions, e.g. Gene Ontology^52^ (GO) biological processes, display similar expression changes. Note that the fold-changes are often very small. (**c**) Precision - recall analysis showing that the treeClust^48,49^ algorithm outperforms three correlation-based coexpression measures. Applying the topological overlap measure (TOM) improves performance further. Annotations in Reactome^47^ were used as gold standard. (**d**) Co-regulation scores for all protein pairs are obtained by combining treeClust with TOM. The score distribution is highly skewed. Where an arbitrary threshold is required, the highest-scoring 0.5% of pairs (N = 62,812) are considered “co-regulated”. (**e**) Co-regulated protein pairs are strongly enriched for subunits of the same protein complex, enzymes catalysing consecutive metabolic reactions and proteins with identical subcellular localization. (**f**) Most proteins are co-regulated with no or few other proteins, but many have more than 5 co-regulated partners. (**g**) Considering proteins that are co-regulated with ≥10 proteins, these groups of co-regulated proteins are almost always enriched in one or more GO terms. (**h**) The global co-regulation map of ProteomeHD created using t-Distributed Stochastic Neighbor Embedding (t-SNE)^56,57^. Distances between proteins indicate how similar their expression patterns are. See www.proteomeHD.net for an interactive version of the map. (**i**) The co-regulation map broadly corresponds to subcellular compartments, and more detailed functional associations can be observed at higher resolution, as exemplified in subpanels 1-3.

### ProteomeHD offers high protein coverage

On average, the 10,323 human proteins in ProteomeHD were quantified on the basis of 28.4 peptides and a sequence coverage of 49% (Supplementary Fig. 1). As expected from shotgun proteomics data, not every protein is quantified in every condition. The 294 input experiments quantify 3,928 proteins on average. Each protein is quantified, on average, in 112 biological conditions (Supplementary Fig. 1). As a rule of thumb, coexpression studies discard transcripts detected in less than half of the samples. However, with 294 conditions ProteomeHD is considerably larger than the typical coexpression analysis. We therefore decided to use a lower arbitrary cut-off and include proteins for downstream analysis if they were quantified in about a third of the conditions. Specifically, we focus our co-regulation analysis on the 5,013 proteins that were quantified in at least 95 of the 294 perturbation experiments. On average, these 5,013 proteins were quantified in 190 conditions; 43% were quantified in more than 200 conditions (Supplementary Fig. 1).

### Machine-learning captures functional protein associations

Proteins that are functioning together have similar patterns of up- and down regulation across the many conditions and samples in ProteomeHD. For example, the patterns of proteins belonging to two well-known biological processes, oxidative phosphorylation and rRNA processing, can be clearly distinguished, even though most expression changes are well below 2-fold (Fig. 1b). Therefore, we reasoned that it should be possible to reveal functional links between proteins on the basis of such regulatory patterns, and reveal the function of unknown proteins by associating them with well-characterized ones.

Traditionally, the extent of coexpression between two genes is determined by correlation analysis, for example using Pearson’s correlation coefficient (PCC). Since PCC is very sensitive to outlier measurements, Spearman’s rank correlation (rho) or Biweight midcorrelation (bicor) are sometimes used as more robust alternatives. We calculated these three correlation coefficients for all 12,562,578 pairwise combinations of the 5,013 protein subset of ProteomeHD. To assess which metric works best for ProteomeHD we performed a precision-recall analysis, using known functional protein - protein associations from Reactome^47^ as gold standard. This showed no major difference between the correlation measures, although Spearman’s rho performs slightly better than the others (Fig. 1c).

We then tested a new type of coexpression measure based on unsupervised machine-learning. Specifically, we used the treeClust algorithm developed by Buttrey and Whitaker, which infers dissimilarities based on decision trees^48,49^. In short, treeClust runs data through a set of decision trees, which it creates without explicitly provided training data, and essentially counts how often two proteins end up in the same leaves. This results in pairwise protein - protein dissimilarities (not clusters of proteins). Importantly, we find that treeClust dissimilarities strongly outperform the three correlation metrics at predicting functional relationships between proteins in ProteomeHD (Fig. 1c).

Finally, we apply a topological overlap measure (TOM)^50,51^ to the treeClust similarities, which further enhances performance by approximately 10% as judged by the area under the precision-recall curve (Fig. 1c). The TOM is typically used to improve the robustness of correlation networks by re-weighting connections between two nodes according to how many shared neighbors they have. The TOM-optimised treeClust results form our “co-regulation score”. This score is continuous and reflects how similar two proteins behave across ProteomeHD, i.e. the higher the score the more strongly co-regulated two proteins are. However, for some questions a simplified categorical interpretation is more straightforward. In these cases we arbitrarily consider the top-scoring 0.5% percent of proteins pairs as “co-regulated”. In this way, we identify 62,812 co-regulated protein pairs (Fig. 1d, Supplementary Table 3). For comparison, if the same data were analysed by Pearson’s correlation, selecting the top 0.5% pairs would correspond to a cut-off of PCC > 0.69, which is generally considered a strong correlation.

We then tested whether co-regulation indicates co-function. Indeed, we find that co-regulated protein pairs are heavily enriched for subunits of the same protein complex, enzymes catalysing consecutive metabolic reactions and proteins occupying the same subcellular compartments (Fig. 1e). The majority of proteins are co-regulated with at least one other protein, and about a third have more than five co-regulation partners (Fig. 1f). For 99% of the tested proteins that had ≥ 10 co-regulated pairs, the group of their co-regulation partners is enriched in at least one Gene Ontology^52^ biological process (Fig. 1g).

### Quantitative protein co-regulation is more informative than co-occurrence

While decision trees are well-understood building blocks of many established machine-learning algorithms, treeClust itself is a relatively recent invention^48^. It was therefore unclear which type of information treeClust captures from a dataset. For example, treeClust scores could simply reflect whether or not two proteins are detected in the same set of samples, a measure that has been successfully exploited previously^41^. To test that we compared treeClust scores to the Jaccard index^53^, a dedicated measure of co-occurrence (Supplementary Fig. 2). In addition, we forced treeClust to learn dissimilarities solely based on co-occurrence by using a “binary” version of ProteomeHD, where all SILAC ratios were turned into ones and all missing values into zeroes. We find that the Jaccard index and “binary” treeClust detect functionally related proteins equally well, but with much lower precision than standard treeClust. This suggests that protein co-regulation, i.e. coordinated changes in protein abundance, rather than co-detection is essential for treeClust performance.

**Figure 2.**
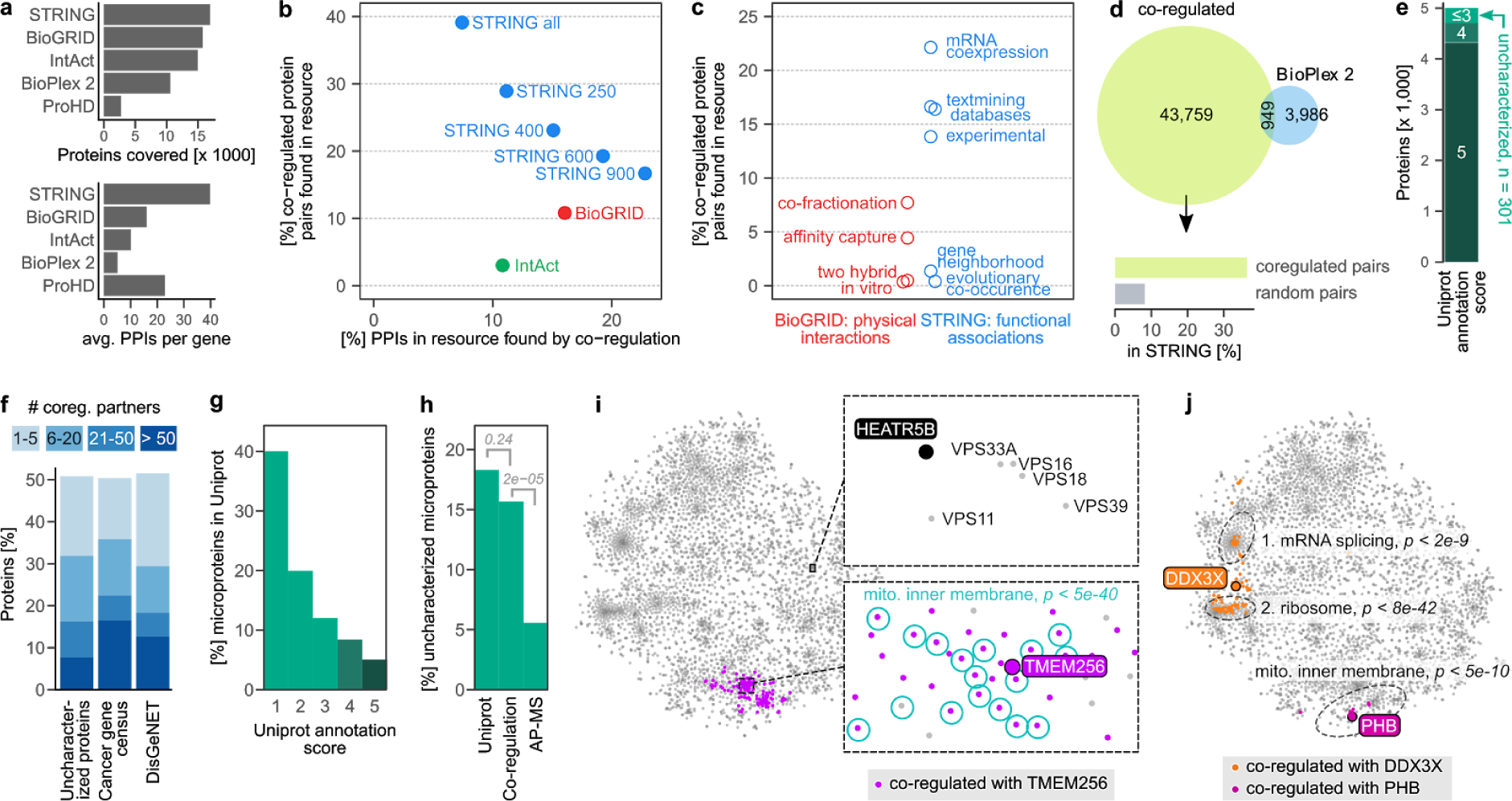
Protein co-regulation complements existing methods and predicts functions of unknown proteins. (**a**) Coverage of protein - protein interactions (PPIs) in comparison to other resources. Top barchart shows the number of genes covered, i.e. having at least one PPI above cut-off. STRING cut-off used: medium (400). Bottom chart shows the average number of PPIs of covered genes. The co-regulation map (ProHD) covers fewer genes than STRING, BioGRID, IntAct and BioPlex 2, but covers many associations between those genes. (**b**) Overlap between PPIs discovered by protein co-regulation and PPIs already present in large-scale annotation resources that cover both physical (BioGrid^60^ and IntAct^59^) and functional (STRING^61^) associations. Multiple association score cut-offs were considered for STRING. These three resources integrate data from many small and large-scale studies. (**c**) Coverage of co-regulated protein pairs in BioGRID and STRING broken down by the type of functional genomics evidence available in each resource. (**d**) Number of co-regulation links compared to PPIs found for the same set of genes by BioPlex 2.0^4^, one of the largest PPI datasets reported to date by a single study. Associations unique to co-regulation are strongly enriched for links in STRING, compared to random gene pairs. (**e**) Out of the 5,013 proteins in the co-regulation map, 301 have a UniProt annotation score ≤3 and are thus defined as uncharacterized. (**f**) Connectivity of either uncharacterized proteins or proteins encoded by disease genes to well-characterized proteins (annotation score ≥4). 51% of uncharacterized proteins have at least one co-regulation partner, 32% have more than five. (**g**) Barchart showing the percentage of all 20,408 human UniProt (SwissProt) proteins that are microproteins, i.e. have a molecular weight < 15 kDa. Note that microproteins are heavily enriched among less well-characterized proteins. (**h**) 18% of uncharacterized proteins in UniProt are microproteins, compared to 16% of the uncharacterized proteins in the co-regulation map and 6% in state-of-the-art AP-MS experiments, represented by BioPlex. *P*-values are from one-sided Fisher’s Exact test. (**i**) The uncharacterized microprotein TMEM256 has many co-regulation partners, which are enriched for GO term “mitochondrial inner membrane” among others. Bonferroni-adjusted *P-*value is from a hypergeometric test. The uncharacterized HEATR5B protein has no co-regulation partners above the default threshold, but its position in the map nevertheless indicates a potential function. (**j**) For multifunctional proteins, co-regulation can reveal a mix of their functions (DDX3X), or their main function only (prohibitin, PHB). Three representative GO terms are shown.

Furthermore, it remained unclear what type of quantitative relationships treeClust can identify and why it outperforms correlation metrics for protein coexpression analysis. We addressed this in a separate study by systematically benchmarking treeClust using synthetic data^54^ (available at: www.biorxiv.org/content/10.1101/578971v1). In short, we found that treeClust detects linear but not non-linear relationships. Unlike correlation metrics, it distinguishes between strong, tight-fitting relationships and weak trends. Finally, as may be expected from an algorithm based on decision trees, it is exceptionally robust against outliers. These properties of treeClust collectively explain its superior performance on ProteomeHD^54^. However, experiments with synthetic data also show that treeClust works best for large datasets with 50 samples or more, depending on additional parameters such as the frequency of missing values. Traditional correlation analysis may be better suited for smaller gene expression datasets^54^.

### A co-regulation map of the human proteome

As a result of treeClust learning we know for each protein how strongly - or weakly - it is co-regulated with any other protein. In principle, these results could be displayed as a scale-free protein interaction network with edges indicating co-regulation (Supplementary Fig. 3). However, due to size and nature of our co-regulation data - 62,812 top-scoring links between 5,013 proteins - it appears impossible to avoid low-informative “hairball” graphs^55^.

**Figure 3.**
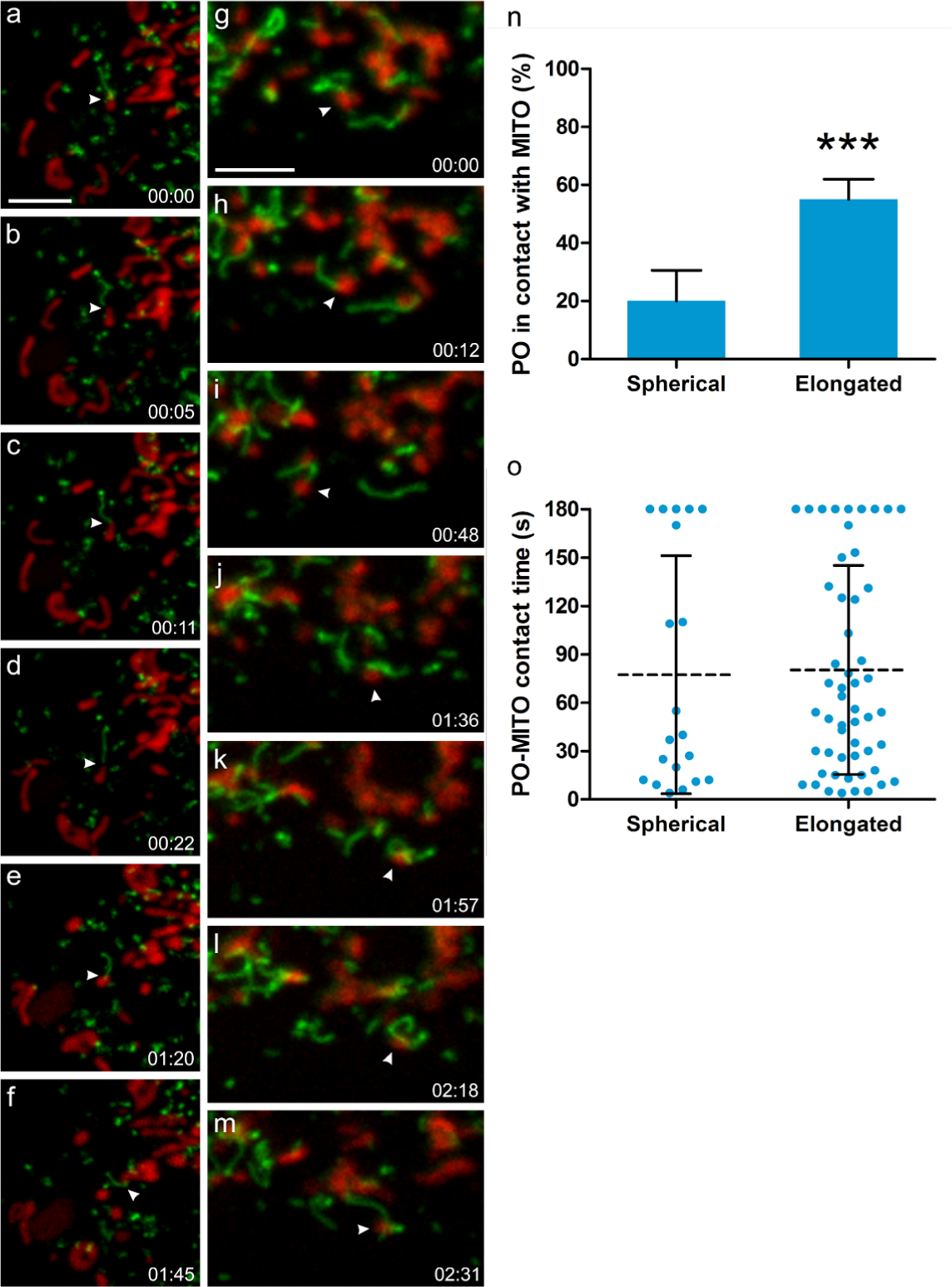
PEX11β mediates the formation of peroxisomal membrane protrusions which interact with mitochondria in mammalian cells. (**a-m**) COS-7 cells were transfected with PEX11β-EGFP, mitochondria were stained with Mitotracker (red) and cells observed live using a spinning disc microscope. PEX11β, a membrane shaping protein, induces the formation of tubular membrane protrusions from globular peroxisomes. We show here that those membrane protrusions can interact with mitochondria. (**a-f**) shows a peroxisome which interacts with a mitochondrion via its membrane protrusion (arrowhead), and follows it, occasionally detaching and re-establishing contact before interacting with another mitochondrion (see Supplemen-tary Movie 1). (**g-m**) shows a mitochondrion (arrowhead) which interacts with a peroxisome via a peroxisomal membrane protrusion. It then detaches and moves away to interact with another peroxisome, which wraps its protrusion around it, before interacting with another mitochondrion (see Supplementary Movie 2). (**n**) Quantification of interactions between spherical or elongated peroxisomes (PO) with mitochondria (MITO). The average result of 3 independent experiments is shown, error bars indicate standard deviation. (**o**) Quantification of contact time. Note that elongated PO interact more frequently with MITO than spherical PO, but for similar time periods. PO-MITO interactions are generally long-lasting (see Supplementary Movie 3) (n=200 peroxisomes from 5 different cells). Dotted line indicates the mean, error bars indicate standard deviation. *** *P* < 0.001 from a two-tailed unpaired *t* test; Time (min:sec). Scale bars, 5 µm.

We therefore chose to visualize the protein - protein co-regulation matrix using t-Distributed Stochastic Neighbor Embedding (t-SNE)^56,57^. This produces a two-dimensional proteome co-regulation map in which the distance between proteins indicates how similar they responded to the various perturbations in ProteomeHD (Fig. 1h, Supplementary Table 4). Notably, t-SNE takes all pairwise co-regulation scores into account, rather than focussing on a small number of links above an arbitrary threshold. The t-SNE map shows that protein co-regulation is closely related to co-function. From a global perspective, the map reflects the subcellular organization of the cell (Fig. 1i). It broadly separates organelles and, for example, sets apart the nucleolus from the nucleus. A closer look into three sections of the map reveals that it captures more detailed functional relationships, too. For example, the five protein complexes of the respiratory chain are almost resolved (Fig. 1i, section 1). The section also contains the phosphate and ADP carriers that transport the substrates for ATP synthesis through the inner mitochondrial membrane, and ATPIF1 - a short-lived, post-transcriptionally controlled key driver of oxidative phosphorylation in mammals^58^. Similarly, cytoskeleton proteins such as actins and myosins are found next to their regulators, including Rho GTPases and the Arp2/3 complex (Fig. 1i, section 2). A third example section shows groups of proteins involved in RNA biology, from nucleolar rRNA processing to mRNA splicing and export (Fig. 1i, section 3). Notably, these annotations are only used to illustrate that the co-regulation map reflects functional similarity; the map itself is generated without any curated information, solely on the basis of protein abundance changes in ProteomeHD. Therefore, the co-regulation map provides a data-driven overview of the proteome, connecting proteins into functionally related groups.

### Co-regulation complements existing functional genomics methods

We next asked if protein co-regulation can predict associations that are not detected by other methods. For this we compare co-regulation to four alternative large-scale resources: IntAct^59^, BioGRID^60^, STRING^61^ and BioPlex^4^. The first three are “meta-resources”, i.e. they compile curated sets of protein - protein interactions (PPIs) from the results of thousands of individual studies. Since meta-resources generally map interactions to gene loci rather than proteins, we disregard protein isoforms for this comparison and focus on co-regulated genes.

The co-regulation map covers fewer distinct genes than the other resources, but only STRING captures more interactions per average gene (Fig. 2a). Based on the 2,565 genes covered by both approaches, around 39% of the gene pairs identified as co-regulated had previously been linked in STRING (Fig. 2b). This suggests that co-regulation analysis confirms existing links, but also provides many additional ones. Conversely, only 7% of STRING PPIs are co-regulated, which may reflect the diverse molecular nature of associations covered by STRING. Notably, the overlap between the resources depends on the stringency setting: considering fewer, more stringent STRING interactions decreases the coverage of co-regulated genes and increases STRING PPIs identified as co-regulated (Fig. 2b). An equivalent trend would be observed when modulating the co-regulation cut-off. STRING associations are based on multiple types of evidence, of which “mRNA coexpression” unsurprisingly shows the highest individual overlap with protein co-regulation results (Fig. 2c).

Next, we compared co-regulation specifically to physical PPIs catalogued by IntAct and BioGRID. We find that 11% of co-regulated gene pairs have a documented physical interaction between their proteins in BioGRID, and 3% are found in the smaller IntAct database (Fig. 2b). These physical PPIs were mainly derived from co-fractionation experiments, which tend to capture indirect interactions, rather than methods that detect direct interactions, such as two-hybrid screens (Fig. 2c).

Finally, we compared the co-regulation approach to an individual functional genomics project: BioPlex 2.0, the most comprehensive affinity purification–mass spectrometry (AP-MS) study reported to date^4^. BioPlex reports 4,935 physical interactions between the proteins used in our study, of which 19% are also co-regulated (Fig. 2d). An additional 43,759 potential links between these proteins are identified uniquely by co-regulation. These are strongly enriched for functional protein associations found in STRING, compared to a random set of protein pairs (Fig. 2d). In conclusion, these comparisons suggest that protein co-regulation identifies protein - protein associations in a way that is reliable yet complementary to existing functional genomics methods. Note that proteins can interact physically or genetically or co-localize without being co-regulated, and vice versa. Therefore, protein co-regulation is complementary not just in terms of identifying new links, but also in providing additional, independent biological evidence for associations detected by other approaches.

### Uncharacterized proteins in ProteomeHD are rich in microproteins

The co-regulation map contains 301 uncharacterized proteins, which we define as proteins with a UniProt^62^ annotation score of 3 or less (Fig. 2e). Of these, 51% are co-regulated with at least one fully characterized protein, i.e. a protein with an annotation score of 4 or 5 (Fig. 2f). On median, these uncharacterized proteins have 9 well-studied co-regulation partners, making it possible to predict their potential function in a “guilt by association” approach. We observe a similar connectivity for the cancer gene census^63^, i.e. genes that cause cancer when mutated, and for DisGeNET^64^ genes, which are genes implicated in a broad range of human diseases (Fig. 2f). Therefore, protein co-regulation may also be helpful for functional analysis of human disease genes.

A common property of uncharacterized proteins is their small size. For example, proteins smaller than 15 kDa constitute 18% of the uncharacterized proteins in the human proteome, but only 5% of the characterized ones. Among the least well understood fraction of the proteome, i.e. proteins with an annotation score of 1, 40% are smaller than 15 kDa (Fig. 2g). This discrepancy is set to increase further, since hundreds or thousands such microproteins have so far been overlooked by genome annotation efforts^65,66^. Microproteins can regulate fundamental biological processes^67^, but their small size makes it difficult to identify interaction partners^65,68^ or to target them in mutagenesis screens^65^. Microprotein sequences also tend to be less conserved than those of longer protein-coding genes^69^. We reasoned that our perturbation proteomics approach may help to reduce the annotation gap for small proteins. As it only requires proteins to be quantifiable in cell extracts we expect it to be less biased by protein size than methods involving extensive genetic or biochemical sample processing. Indeed, we find that 16% of the uncharacterized proteins in the co-regulation map are smaller than 15 kDa, which is close to the 18% in the proteome overall (Fig. 2h). However, it is a significant difference to BioPlex’s cutting-edge AP-MS data, in which microproteins drop to 6% (*p* < 2e-5 in a one-tailed Fisher’s Exact test).

The fact that microproteins are not underrepresented in ProteomeHD does not automatically mean that their detection and characterisation is as robust as that of larger proteins. However, the average microprotein in the co-regulation map has been identified by 12.2 peptides, many of which overlap and together result in an average sequence coverage of 76.4% (Supplementary Fig. 4a, d). While in a typical SILAC experiment proteins are considered to be quantifiable from upwards of two independent observations (SILAC ratio counts), microproteins in the co-regulation map are quantified with an average of 9 ratio counts per experiment, totalling a median of 671 ratio counts across ProteomeHD (Supplementary Fig. 4b, c). This indicates that microprotein quantitation in ProteomeHD is robust. Surprisingly, we find that microproteins have more co-regulation partners than larger proteins, and the same is true for their connectivity in STRING (Supplementary Fig. 4f). Within STRING, the majority of microprotein interactions are derived from curated annotations rather than high-throughput efforts such as RNA coexpression and text mining (Supplementary Fig. 4g). Note that, based on BioGRID, microproteins engage in fewer physical PPIs than larger proteins. This may be the result of an experimental bias (microproteins may dissociate more easily during purification and are more difficult to detect) or reflect a biological property (microproteins may have fewer physical interaction partners). In either case, co-regulation offers itself as a powerful alternative approach to study microprotein functions in a systematic way.

**Figure 4.**
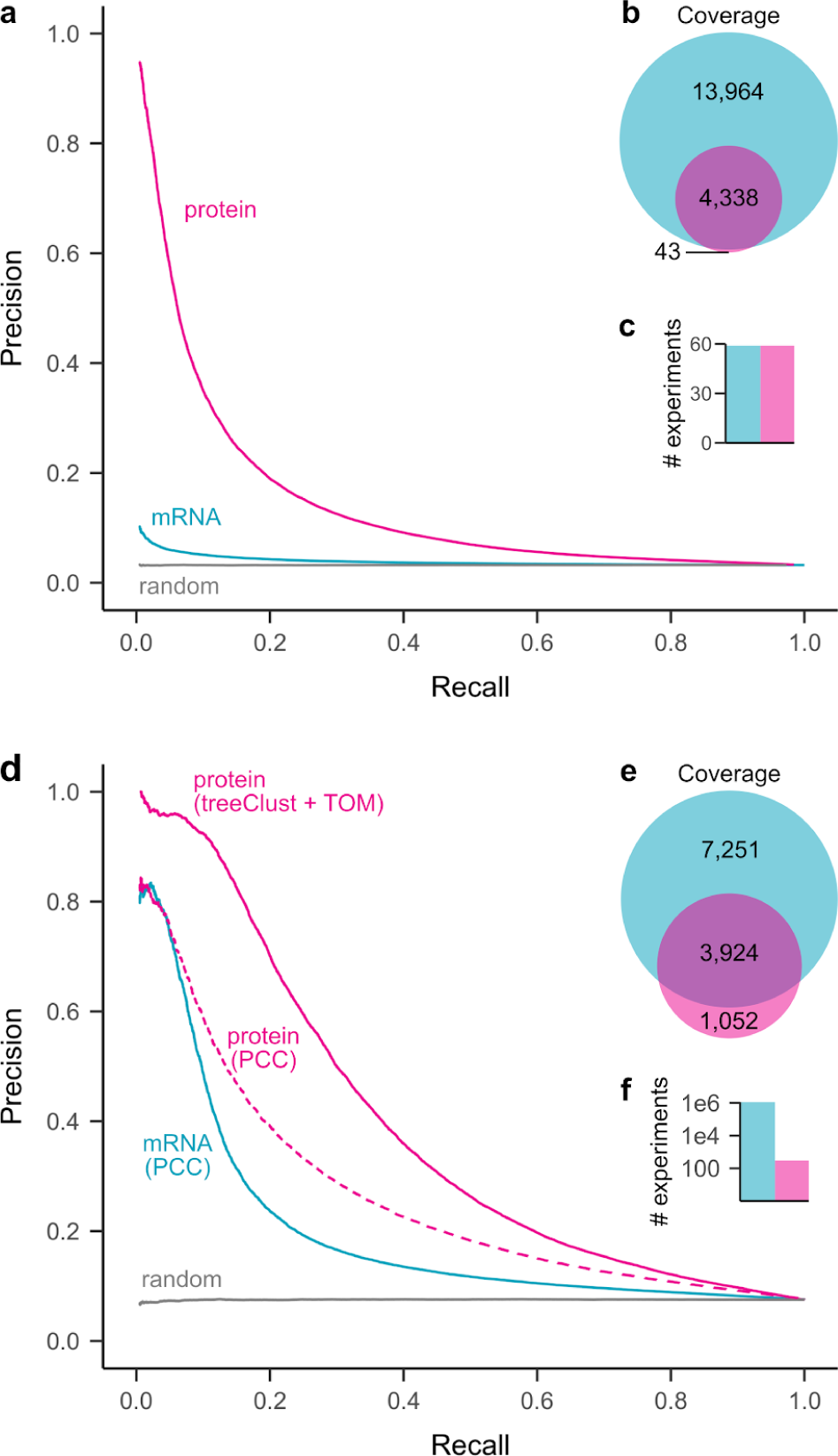
Protein co-regulation enables higher precision from less data, but has lower coverage than classic mRNA coexpression. (**a**) Precision-recall analysis of treeClust machine-learning on a subset of ProteomeHD, that is 59 samples for which matching RNA-seq data were available from a separate study^86^. Reactome pathways were used as gold standard for true functional associations (proteins found in same pathway) and false associations (never found in same pathway). Only annotated genes covered by both datasets were considered for PR analysis (n = 2,901). (**b**) Venn diagram showing number of genes covered by each analysis. (**c**) Barchart showing number of experiments the curves are based on. (**d**) Similar precision-recall analysis of treeClust machine-learning on the full ProteomeHD database, in comparison to Pearson correlation obtained by STRING^61^ on the basis of one million human mRNA profiling samples deposited in the NCBI Gene Expression Omnibus^87^ (“mRNA / PCC”). Protein co-regulation outperforms mRNA correlation despite being based on orders-of-magnitude less data. This is partially due to the use of machine-learning, as predicting associations from ProteomeHD using PCC decreases performance markably (“protein / PCC”). Only annotated genes covered by both datasets were considered for the PR analysis (n = 2,743). (**e, f**) same as (b, c).

### Functional annotation of proteins by co-regulation

To facilitate the characterization of proteins through co-regulation we created the website www.proteomeHD.net. It allows users to search for a protein of interest, showing its position in the co-regulation map together with any co-regulation partners (Supplementary Fig. 5). The online map is interactive and zoomable, making it easy to explore the neighborhood of a query protein. The co-regulation score cut-off can be adjusted and statistical enrichment of Gene Ontology^52^ terms among the co-regulated proteins is automatically calculated.

For example, protein co-regulation can be used to predict the potential function of uncharacterized microproteins such as the mitochondrial proteolipid MP68. MP68 is co-regulated with subunits of the ATP synthase complex, suggesting a function in ATP production (Fig. 1i, section 1). Despite being only 6.8 kDa small, its presence in the co-regulation map is documented by 8 distinct peptides that were observed a total of 398 times across 142 experiments (Supplementary Fig. 4e). Intriguingly, MP68 co-purifies biochemically with the ATP synthase complex, but only in buffers containing specific phospholipids^70,71^, and knockdown of MP68 decreases ATP synthesis in HeLa cells^72^.

Virtually nothing is known about the 12 kDa microprotein TMEM256, although sequence analysis suggests it may be a membrane protein. Its position in the co-regulation map (Fig. 2i) and GO analysis of its co-regulation partners indicates that it likely localizes to the inner mitochondrial membrane (GO:0005743, Bonferroni adj. *p* < 5e-40), where it may participate in oxidative phosphorylation (GO:0006119, *p* < 3e-35).

Some proteins have no co-regulation partners above the default score cut-off, but can still be functionally annotated through the co-regulation map. The uncharacterized 224 kDa protein HEATR5B, for example, is located in an area related to vesicle biology (Fig. 2i). Its immediate neighbours are five subunits of the HOPS complex, which mediates the fusion of late endosome to lysosomes. The position in the map shows that the HOPS complex is the closest fit to HEATR5B’s regulation pattern, but they are not as similar as the top-scoring pairs in our overall analysis. If the co-regulation score cut-off is lowered, HOPS subunits and other endolysosomal proteins are eventually identified as co-regulated with HEATR5B, with concomitant enrichment of the related GO terms. This suggests that HEATR5B may not itself be a HOPS subunit, but could have a related vesicle-based function. Notably, a biochemical fractionation profiling approach also predicted HEATR5B to be a vesicle protein^73^.

Multifunctional proteins appear to fall into two categories in terms of co-regulation behavior. Prohibitin, for example, functions both as a mitochondrial scaffold protein and a nuclear transcription factor^74^. However, only the mitochondrial function is represented in the co-regulation map (Fig. 2j). This could indicate that its nuclear activity is not relevant in the biological conditions covered by ProteomeHD, or that only a small intracellular pool of prohibitin is nuclear, so that changes in its nuclear abundance are insignificant in comparison to the mitochondrial pool. In contrast, the helicase DDX3X shuttles between nucleus and cytoplasm, functioning both as nuclear mRNA processing factor and cytoplasmic regulator of translation^75^. In the co-regulation map, DDX3X sits between the areas related to these two activities and is significantly co-regulated both with proteins involved in nuclear RNA biology and with translation factors (Fig. 2j). Therefore, DDX3X is a multifunctional protein whose separate activities result in a mixed regulatory pattern.

The protein co-regulation data presented here have been integrated into the recently released 11th version of STRING^76^ (https://string-db.org/). In STRING’s human protein - protein association network, links between proteins inferred from co-regulation in ProteomeHD are shown as network edges of the “coexpression” type (Supplementary Fig. 6). Therefore, STRING is an alternative source for users wishing to explore protein co-regulation in conjunction with other types of association evidence.

### A new function for PEX11β in peroxisome-mitochondria interplay

Some well-characterized proteins have unexpected co-regulation partners. For example, PEX11β is a key regulator of peroxisomal membrane dynamics and division^77^. However, PEX11β’s co-regulation partners are not peroxisomal proteins but subunits of the mitochondrial ATP synthase and other components of the electron transport chain (Fig. 1i, section 1). These proteins are located to the inner mitochondrial membrane, making a physical interaction with PEX11β unlikely. However, peroxisomes and mitochondria in mammals are intimately linked cooperating in fatty acid β-oxidation and ROS homeostasis^78^. How these organelles communicate or mediate metabolite flux has been elusive. Live cell imaging revealed that expression of PEX11β-EGFP in mammalian cells induced the formation of peroxisomal membrane protrusions, which interact with mitochondria (Fig. 3, Supplementary movies 1-3). Interactions of elongated peroxisomes with mitochondria were more frequent than those of spherical organelles, but both interactions were long-lasting (Fig. 3n,o). This indicates that peroxisome elongation can facilitate organelle interaction, but once organelles are tethered, the duration of contacts is similar between different morphological forms. Miro1 (RHOT1), a membrane adaptor for the microtubule-dependent motors kinesin and dynein^79^, is also co-regulated with PEX11β (Fig. 1i, section 1). We and others recently showed that Miro1 distributes to mitochondria and peroxisomes^80,81^ indicating that it coordinates mitochondrial and peroxisomal dynamics with local energy turnover. Peroxisome-targeted Miro1 (Myc-Miro-PO) can be used as a tool to exert pulling forces at peroxisomal membranes, which results in the formation of membrane protrusions in certain cell types^82^ (Supplementary Fig. 7). We show here that silencing of PEX11β inhibits membrane elongation by Myc-Miro-PO, confirming that PEX11β is required for the formation of peroxisomal membrane protrusions (Supplementary Fig. 7). These findings are in agreement with studies in plants, where *At*PEX11a has been reported to mediate the formation of peroxisomal membrane extensions in response to ROS^83^. In yeast, peroxisome-mitochondria contact sites are established by *Sc*Pex11 and *Sc*Mdm34, a component of the ERMES complex^84^. Additional tethering functions for the yeast mitofusin Fzo1 and ScPex34 in peroxisome–mitochondria contacts have recently been revealed^85^. Importantly, the study also demonstrated a physiological role for peroxisome–mitochondria contact sites in linking peroxisomal β-oxidation and mitochondrial ATP generation by the citric acid cycle^85^. We conclude that PEX11β and Miro1 contribute to peroxisome membrane protrusions, which present a new mechanism of interaction between peroxisomes and mitochondria in mammals. They likely function in the metabolic cooperation and crosstalk between both organelles, and may facilitate transfer of metabolites such as acetyl-CoA and/or ROS homeostasis during mitochondrial ATP production. These findings now enable future studies on the precise functions of peroxisome membrane protrusions in mammalian cells and the role of PEX11β.

### Proteomics enables higher accuracy but lower coverage than transcriptomics

To compare the impact of mRNA and protein abundances on expression profiling we first focussed on 59 SILAC ratios in ProteomeHD that measured abundance changes across a panel of lymphoblastoid cell lines^30^. For these samples, corresponding mRNA abundance changes have been determined using RNA-sequencing^86^. Repeating treeClust learning on the basis of these data, we observed that protein coexpression predicts functional associations with far higher precision than mRNA coexpression (Fig. 4a). Similar results have recently been reported for a panel of human cancer samples^19^.

Such analyses show that in a direct gene-by-gene, sample-by-sample comparison, protein expression levels are better indicators for gene function than mRNA expression. However, the amount of transcriptomics data published to date vastly exceeds that of proteomics studies. For example, the NCBI GEO repository currently holds mRNA expression profiling data from more than one million human samples^87^. This raises the possibility that the sheer quantity of available transcriptomics data could overcome their reduced reflection of functional links and, in combined form, perform better than protein-based measurements. To test this we compared the ProteomeHD co-regulation score with Pearson correlation coefficients obtained by STRING, which leverages the vast amount of mRNA expression experiments deposited in GEO^61,88^. Remarkably, precision-recall analysis shows that the protein co-regulation score still outperforms mRNA coexpression, despite being based on only 294 SILAC ratios (Fig. 4b). Much of this improvement is due to the robustness of treeClust machine-learning, as Pearson’s correlation coefficients derived from the same ProteomeHD data work only moderately better than mRNA correlation (Fig. 4b). While only gene pairs with both mRNA and protein expression measurements were considered for the precision-recall analysis, the transcriptomics and proteomics datasets individually covered 17,436 and 4,976 genes, respectively (Fig. 4b). Therefore, mRNA profiling outperforms protein profiling in terms of gene coverage. In addition, transcriptomics remains the only expression profiling approach suitable for non-coding RNAs.

## DISCUSSION

ProteomeHD in conjunction with machine learning provides an entry point for “big-data”-type protein co-regulation analysis into the functional genomics methods repertoire. It is possible that accuracy and coverage could be increased further by adding additional proteomics data. To test this we randomly removed 5%, 10% or 15% of the data points in ProteomeHD. This decreases performance reproducibly and proportionally to the amount of removed data (Supplementary Fig. 8), suggesting that ProteomeHD has not reached saturation and expanding it will further enhance its performance. One possibility would be to incorporate other types of proteomics experiments, such as affinity-purifications or indeed the entire PRIDE^46^ repository. The latter approach is for instance taken by the Tabloid Proteome, which infers protein associations based on detecting them in the same subset of many different proteomics experiments^41^. However, there is a benefit of restricting ProteomeHD to perturbation experiments. It supports a biological interpretation of protein associations derived from it: two co-regulated proteins are part of the same cellular response to changing biological conditions, even though the precise molecular nature of the connection remains unknown. In this way, protein co-regulation analysis is analogous to genetic interaction screening. This also sets protein co-regulation apart from indiscriminate protein covariation or co-occurrence analyses, which find protein links in a mix of proteomics data and therefore give no insight into the possible biological connection.

A key difference between our approach and previous gene coexpression studies is our application of two machine-learning algorithms, treeClust^48^ and t-SNE^56,57^. Inferring protein associations through treeClust learning is both more robust and sensitive than a traditional correlation-based approach, providing a leap in the accuracy with which functionally relevant interactions can be identified from the same dataset. For example, a recent study reported a protein co-regulation network across 41 cancer cell lines and subsequently identified dysregulated protein associations that predict drug sensitivities of these cell lines^20^. Applying Spearman’s correlation to high-quality, TMT-based proteomics data allowed Lapek *et al*^20^ to detect protein-protein associations with an accuracy that was tenfold higher than that based on matching mRNA coexpression data. When applying treeClust to these data, strikingly, we can further improve this performance (Supplementary Fig. 9a). This suggests that treeClust may be helpful for the detection of “dysregulation biomarkers” in the future. The second machine-learning tool we apply here, t-SNE, visualizes treeClust-learned protein associations as a 2D map. Correlation networks are typically built from a limited number of the strongest pairwise interactions, whereas t-SNE takes into account the similarity - or dissimilarity - between all possible pairwise protein combinations. It creates the map that best reflects both direct and indirect relationships between all proteins. In this way, also proteins that are not directly linked to the core network can be placed into a functional context. For example, a t-SNE co-regulation map obtained for Lapek *et al*’s cancer proteomics dataset contains the complete set of ∽6,800 proteins, rather than the 3,024 proteins that are directly correlated with another protein (Supplementary Fig. 9b). Moreover, protein-protein associations visualized by t-SNE can be explored in a hierarchical manner, with larger distances indicating weaker co-regulation. This may be useful for studying connections between related protein complexes (Fig. 1i) or to reveal broad functional clues for uncharacterized proteins for which no detailed predictions are available, such as the HEATR5B protein assigned to the vesicle area of the co-regulation map (Fig. 2i). Our web application at www.proteomeHD.net is designed to support researchers in exploring co-regulation data at multiple scales, to validate existing hypotheses or create new ones.

Protein coexpression analysis identifies functional connections between proteins with an accuracy and sensitivity that is substantially higher than traditional mRNA coexpression analysis. This may be particularly important for constitutively active genes, which constitute about half of human genes^44^ and are primarily controlled at the protein level^89,90^. With an ever increasing amount of protein expression data making their way into the public domain, and the simplicity of exploiting the analysis results by the scientific community, protein coexpression analysis has a large potential for gene function annotation. Only 300 quantitative proteomics measurements sufficed in conjunction with machine learning to establish functional connections between many human genes, which may be of considerable interest for proteome annotation in less studied or difficult to study organisms.

## ACKNOWLEDGEMENTS

We are grateful to Damian Szklarczyk for providing the mRNA Pearson correlation data used by STRING and the STRING team for testing our coregulation data and adding it as novel evidence type to STRING 11. We also thank Karen Wills, Kyosuke Nakamura, Constance Alabert and Anja Groth for contributing chromatin enrichment experiments, and Afsoon S. Azadi for support with live-cell-imaging. This work was supported by the Wellcome Trust through a Senior Research Fellowship to J.R. (grant number 103139) and by the Biotechnology and Biological Sciences Research Council (BB/N01541X/1, BB/R016844/1; to M.S.) and H2020-MSCA-ITN-2018 812968 PERICO (to M.S.). The Wellcome Centre for Cell Biology is supported by core funding from the Wellcome Trust (grant number 203149).

## AUTHOR CONTRIBUTIONS

G. K. and J. R. conceived the project. G. K. and P.G. conducted the data analysis. P. G. created the web application. T. A. S., J. B. P. and M. S. conducted the Pex11β analysis. All authors contributed to writing the manuscript.

## SUPPLEMENTARY MOVIE LEGENDS

**Supplementary Movie 1. Interaction of peroxisomal membrane protrusions with mitochondria in COS-7 cells.** *See* Fig. 4a-f.

COS-7 cells were transfected with PEX11β-EGFP, mitochondria were stained with Mitotracker (red), and analysed by live-cell imaging using an IX81 microscope (Olympus) equipped with a CSUX1 spinning disk head (Yokogawa). A peroxisome interacts with a mitochondrion via its membrane protrusion, and follows it, occasionally detaching and re-establishing contact. 200 stacks of 9 planes (0.5 µm thickness, 100 ms exposure) were taken in a continuous stream. 118 frames, 14× speed. Scale bar, 5 µm.

**Supplementary Movie 2. Interaction of peroxisomal membrane protrusions with mitochondria in COS-7 cells.** *See* Fig. 4g-m and legend Movie 1.

Note a peroxisome at the bottom, which interacts with a mitochondrion via its membrane protrusion and then wraps around it, possibly to increase the membrane contact area. 200 stacks of 9 planes (0.5 µm thickness, 100 ms exposure) were taken in a continuous stream. 200 frames, 14× speed. Scale bar, 5 µm.

**Supplementary Movie 3. Interaction of peroxisomal membrane protrusions with mitochondria in COS-7 cells.** *See* legend Movie 1.

A mitochondrion, which moves to the left, is dragging a peroxisome with a membrane protrusion with it, indicating that the organelles are tightly tethered to each other. 200 stacks of 9 planes (0.5 µm thickness, 100 ms exposure) were taken in a continuous stream. 100 frames, 14× speed. Scale bar, 5 µm.

## ONLINE METHODS

### General data analysis and code availability

Data analysis was performed in R^91^. R scripts and input files required to reproduce the results of this manuscript are available in the following GitHub repository: https://github.com/Rappsilber-Laboratory/ProteomeHD. The R package data.table^92^ was used for fast data processing. Figures were prepared using ggplot2^93^, gridExtra^94^, cowplot^95^ and viridis^96^.

### Data selection for ProteomeHD

MS raw data were produced in-house or downloaded from the PRIDE repository^46^. Only experiments fulfilling the following inclusion criteria were considered:

(1) Comparative proteomics experiments, i.e. relative protein quantitations of two or more biological states. For example, cells treated with an inhibitor *vs.* mock control. (2) Biological - not biochemical - comparisons, i.e. fold-changes must have been brought about *in vivo*, not by differential biochemical purification. For example, SILAC-labelled cells were treated with inhibitor or mock control, harvested and combined, and chromatin was enriched on the combined sample. In such cases any observed fold-change reflects the response to the inhibitor in the living cell, for example a protein re-localising from cytoplasm onto chromatin. We did not consider experiments that compared, for example, a whole-cell lysate with a chromatin-enriched fraction, as this would measure the impact of the biochemical enrichment rather than a biological event. (3) Quantitation by “stable isotope labeling by amino acids in cell culture” (SILAC)^45^. (4) Samples of human origin.

In addition to these conceptual considerations, the following restrictions were imposed by the data processing pipeline: (5) The SILAC mass shift introduced by heavy arginine must be distinct from heavy lysine. (6) Raw data acquired on an Orbitrap mass spectrometer. (7) Samples alkylated with iodoacetamide, resulting in carbamidomethylation of cysteines.

In total, we considered 294 experiments (SILAC ratios) from 31 projects. A full list of these is provided in Supplementary Table 2, which also includes the PRIDE identifiers of all previously published datasets.

### In-house data collection

80 experiments were performed in-house and analyzed chromatin-enriched samples. Of these, 65 measured the effect of growth factors, radiation and other perturbations on interphase chromatin, which was prepared using Chromatin Enrichment for Proteomics (ChEP)^97^. About half of these experiments had previously been published^36^. Another 15 experiments documented perturbations specifically on freshly replicated chromatin, which was prepared using Nascent Chromatin Capture (NCC)^98^. All mass spectrometry raw files generated in-house have been deposited to the ProteomeXchange Consortium (http://proteomecentral.proteomexchange.org) via the PRIDE partner repository^46^ with the dataset identifier PXD008888 (this repository will be made public upon acceptance of the manuscript).

### MS raw data processing

The 5,288 MS raw files were processed using MaxQuant 1.5.2.8^99^ on a Dell PowerEdge R920 server. The following default MaxQuant search parameters were used: MS1 tolerance for the first Andromeda search: 20 ppm, MS1 tolerance for the main Andromeda search: 4.5 ppm, FTMS MS2 match tolerance: 20 ppm, ITMS MS2 match tolerance: 0.5 Da, Variable modifications: acetylation of protein N-termini, oxidation of methionine, Fixed modifications: carbamidomethylation of cysteine, Decoy mode set to reverse, Minimum peptide length: 7 and Max missed cleavages set to 2. The following non-default settings were used: In group-specific parameters, match type was set to “No matching”. In global parameters, “Re-quantify” was enabled, minimum ratio count was set to 1 and “Discard unmodified counterpart peptide” was disabled. Also in global parameters, writing of large tables was disabled. SILAC labels were set as group-specific parameters as indicated in Supplementary Table 2. Canonical and isoform protein sequences were downloaded from UniProt^62^ on 28th May 2015, considering only reviewed SwissProt entries that were part of the human proteome. Unprocessed MaxQuant result tables, including peptide evidence data, have been deposited into the PRIDE repository PXD008888.

Protein fold-changes were extracted from the MaxQuant proteinGroups file returned by MaxQuant. Non-normalized SILAC ratios were considered for downstream analysis, log2 transformed and median-normalised. From triple labelling experiments, the heavy/light and medium/light ratios - but not the heavy/medium ratios - were considered. Proteins detected in less than 4 experiments were discarded, as were proteins labeled as contaminants, reverse hits and those only identified by a modification site. The resulting data matrix, ProteomeHD, can be downloaded as Supplementary Table 1.

### Calculation of treeClust dissimilarities

It is common in gene coexpression studies to remove genes that were detected in less than half of the samples from the analysis. However, given the unusually large size of ProteomeHD we chose a different arbitrary cut-off, excluding proteins that were detected in less than 95 (about a third) of the 294 experiments. For the remaining 5,013 proteins in ProteomeHD we used the treeClust^48^ R package to calculate all 12,562,578 pairwise dissimilarities. Note that treeClust was designed not only to measure inter-point dissimilarities but also to perform clustering^48,49^. However, in this study we use it only to calculate dissimilarities, via the treeClust.dist function. The dissimilarity specifier was set to d.num = 2, so that dissimilarities are weighted according to tree quality. We optimised two hyperparameters of treeClust and rpart, which is the routine treeClust uses to create decision trees. These were treeClust’s serule argument, which defines to extent to which trees are pruned, and rpart’s complexity (cp) parameter, which describes the improved fit required to attempt a split. A grid search was performed against the Reactome gold standard (see below) and the area under precision - recall curves was used to identify optimal parameter settings. They were determined to be serule = 1.8 and cp = 0.105, providing approximately a 10% performance improvement over treeClust’s default settings.

### Protein co-regulation scores

To calculate the final pairwise co-regulation scores, treeClust dissimilarities were transformed further. First, they were turned into similarities, i.e. 1 - treeClust dissimilarity. Using the WGCNA^100,101^ R package, we then performed a sigmoid transformation of these treeClust similarities, creating an adjacency matrix. The settings of parameters mu and alpha for this transformation were optimised in a grid search against the Reactome gold standard, using the area under precision - recall curves as readout. In a third step, the adjacency matrix was transformed into a topological overlap matrix using WGCNA’s TOMsimilarity function, with the TOMDenom parameter set to “mean”. These TOM similarities are the co-regulation scores used throughout our analysis. Co-regulation scores for all of our 12,562,578 protein pairs can be downloaded from the PRIDE repository PXD008888.

While the co-regulation score is continuous, some analyses benefitted from a simplified categorical approach. For these cases we arbitrarily defined the highest-scoring 0.5% of protein pairs as “co-regulated pairs” and the remaining 99.5% of pairs as “not co-regulated pairs”. A list of all 62,812 co-regulated protein pairs is available as Supplementary Table 3.

### Reactome gold standard

A gold standard set of reference proteins was defined using Reactome^47^. Bona fide functionally associated protein pairs (true positives) were defined as protein pairs found in the same “detailed” Reactome pathway. This was inferred from the file UniProt2Reactome.txt (available at https://reactome.org/download-data), where each protein is annotated to the lowest level subset of Reactome pathways. To make sure that only closely related protein pairs were assigned the “true positive” label, we excluded two pathways that were composed of > 200 proteins. We defined protein pairs that are not functionally associated (false positives) as proteins that are never in the same Reactome pathway, at any annotation level. This was inferred from UniProt2Reactome_All_Levels.txt (also available at https://reactome.org/download-data), a file that maps proteins to all levels of the Reactome pathway hierarchy. A copy of this gold standard is available in the Github repository noted above.

### Comparison of treeClust and correlation metrics

Pearson’s correlation coefficients (PCC) and Spearman’s rank correlation coefficients (rho) were obtained using the cor function in R, for the same protein pairs covered by the treeClust analysis. Biweight mid-correlation coefficients (bicor) were calculated with default settings using the R package WGCNA^101,102^. Changing the maxPOutliers parameter of the bicor function did not improve performance. Precision - recall (PR) analysis was performed with the ROCR package^103^ using true and false positive pairs compiled from annotation in Reactome (see paragraph Reactome gold standard). The random classifier was created by scrambling co-regulation scores.

### t-SNE visualization

To visualize ProteomeHD as a 2D co-regulation map, co-regulation scores were subjected to t-Distributed Stochastic Neighbor Embedding (t-SNE)^56,57^ using the Rtsne^104^ package for R. The theta parameter was set to zero to calculate the exact embedding. The perplexity parameter was set to 50, up from the default of 30, to account for the large size of the co-regulation dataset. 1,500 iterations were performed. However, visual comparison of the t-SNE maps showed that these parameter adaptations provided only a marginal improvement over the default settings. Organelles were labelled based on subcellular locations assigned by UniProt^62^ to these proteins, zoom regions were annotated manually based on available literature. Plot coordinates and annotations are available as Supplementary Table 4.

### Network visualizations

In addition to t-SNE, the protein co-regulation matrix was also visualized as an undirected, weighted network using the igraph^105^ and GGally^106^ packages in R. The network contains the same 5,013 proteins as the co-regulation map, but only considers links above the arbitrary co-regulation threshold, i.e. between the top-scoring 0.5% of protein pairs. For these pairs, the network edges are weighted by the co-regulation score. A set of common network layout algorithms were deployed through the sna (social network analysis)^107^ R package.

### Testing for co-functionality among of co-regulated proteins

To test if protein co-regulation reflects co-function we defined three sets of “functionally related” protein pairs (subunits of the same protein complexes, enzymes catalyzing consecutive metabolic reactions and proteins with identical subcellular localization) as previously described^25^.

To test larger groups (not pairs) of co-regulated proteins for functional enrichment, we analyzed enrichment of Gene Ontology terms using the topGO^108^ R package. For each protein we tested the group of its co-regulation partners for GO term enrichment. Because some proteins are co-regulated with no or very few other proteins, we restricted the analysis to proteins that are co-regulated with at least 10 proteins. The three aspects (Biological process, Molecular function, Cellular component) of GO were downloaded from QuickGO^109^ with taxon set to human and qualifier to null. Rather than the whole proteome, only proteins that were included in the treeClust analysis and had GO annotations were used as the gene “universe” or background for the topGO analysis. Enrichment of GO terms among protein co-regulation groups was tested considering GO graph structure and using a Fisher’s exact test.

### Annotation of the co-regulation map

Proteins localizing to specific subcellular compartments were downloaded from UniProt^62^ using the following tags: Nucleus (SL-0191), Nucleolus (SL-0188), Endoplasmic reticulum (SL-0095), Mitochondrion (SL-0173), Cytoplasm (SL-0086), Secreted (SL-0243). Proteins and protein complexes in zoom regions (Fig. 1i) were annotated individually based on the available literature.

### Creating the www.proteomeHD.net framework

The ProteomeHD online application was written in Python Flask web framework. The interactive plots are generated using Bokeh visualization library for Python (https://github.com/bokeh/bokeh). The Gene Ontology and KEGG enrichment statistics are obtained from a STRING^61^ server using an API call with maximally top 100 proteins co-regulated with the query. Only significantly enriched terms (hypergeometric test, Bonferroni adjusted *P* value < 0.1) are displayed.

### Comparison to orthogonal methods

Physical protein-protein-interactions (PPIs) detected by a comprehensive range of small- and large-scale methods were assessed using BioGRID^60^, version 3.4.152. Data from IntAct^59^ were used as a smaller but curated resource of physical PPIs. Functional protein associations mapped by a large range of methods and publications were inferred from STRING^61^, version 10.5. Note that the protein co-regulation scores described here are only used by STRING starting with version 11^76^. BioPlex 2.0^4^ served as an example for physical interactions mapped by a single project.

### Annotation of uncharacterized and disease genes

Proteins were defined as “uncharacterized” on the basis of having an annotation score ≤ 3 in UniProt^62^. The UniProt annotation score is a heuristic measure of the annotation state of a protein, expressed as a 5-point system (www.uniprot.org/help/annotation_score). The score combines various types and layers of UniProt annotation, and weights manually curated evidence higher than automated annotation. It may not always agree with the state of “characterization” that field experts would assign to the same protein. However, as an unbiased, data-driven approach we believe the UniProt annotation score is better suited to systematically identify uncharacterized proteins than manual annotation could be. Even with a systematic way of measuring the degree of annotation, the definition of what constitutes an “uncharacterised” protein is an arbitrary one. We chose “3 points or less” as the “uncharacterized” cut-off, because the available information for such proteins tends to be very vague, e.g. a sequence-based prediction as “multi-pass membrane protein”. In contrast, we found that the biological function of most 4-star proteins could be established reasonably well from the available literature.

The Cancer Gene Census, i.e. genes that can cause cancer when mutated, was curated by COSMIC (Catalogue Of Somatic Mutations In Cancer, version 81)^63^. DisGeNET was used as a comprehensive, curated list of human gene - disease associations^64^.

### Comparison of mRNA and protein expression profiling

For the comparison of matched samples and proteins we considered mRNA and protein expression changes across 59 lymphoblastoid cell lines (Fig. 4a). The protein fold-changes are part of ProteomeHD and were originally published by Battle and colleagues^30^. RNA-sequencing data for the same cell lines and proteins were also previously reported^86^. We used the RNA-sequencing data to calculate mRNA fold-changes relative to a 60th cell line, which was the same cell line used as a SILAC reference for the protein expression data. The combined mRNA and protein dataset has been described in more detail elsewhere^25^. Fold-changes for genes covered by both the transcriptomics and proteomics analysis were subjected to treeClust learning (default parameters) and PR curves were obtained as described above.

For a more comprehensive comparison we considered protein associations predicted using treeClust learning or PCC on the basis of all 294 SILAC ratios in ProteomeHD (Fig. 4b). This was compared to mRNA associations inferred by PCC on the basis of all human mRNA expression data processed by STRING. STRING’s state-of-the-art mRNA coexpression analysis pipeline considers all microarray and RNA-sequencing data deposited in the GEO repository^87^, resulting in one of the largest mRNA coexpression analyses available to date^61,88^. Note that for this comparison we did not use the STRING coexpression score, which is calibrated against the KEGG database, but the original uncalibrated Pearson’s correlations, which were kindly provided by Damian Szklarczyk. STRING PCCs are calculated separately for one- and two-channel microarrays and RNA-sequencing experiments. We used the average of these for the precision - recall analysis, which performed better than any individual experiment type.

### Validation of treeClust and t-SNE on the cancer proteomics dataset

Lapek *et al* measured the abundances for 6,911 proteins in 41 different breast cancer cell lines^20^. These data are available as Supplementary Table 2 (tab 3) of their report. As described by Lapek *et al*, we converted the protein intensities into log2 fold-changes over the median intensity measured for each protein across all cell lines. We then calculated Pearson’s, Spearman’s rank and bicor correlations for all possible protein pairs, as for ProteomeHD. The Spearman’s correlation coefficients obtained in this way are identical to the ones obtained by Lapek *et al* using the cor.prob function (Supplementary Table 6 in their report^20^). We also determined treeClust co-regulation scores for all protein pairs. However, treeClust can only grow one decision tree per input variable, i.e. 41 in this dataset, which would be too few for it to perform properly. To circumvent this, we forced treeClust to generate 1,000 decision trees by applying it iteratively. We created 100 treeClust forests, each generated with a random subset of 10 of the 41 variables, and used the average co-regulation score for downstream analysis. Precision-recall analysis using a Reactome gold standard and t-SNE visualization were performed as described above. The CORUM protein complexes displayed in Lapek *et al*’s Figure 2, reported in their Supplementary Table 7^20^, were color-coded in the co-regulation map.

### Comparison of protein co-regulation and co-occurrence

Two different approaches were used to measure protein co-occurrence in ProteomeHD. First, the Jaccard / Tanimoto similarity coefficient^53^ was calculated using the Jaccard package for R. Second, a binary version of ProteomeHD was created, where all SILAC ratios were represented by 1s (“protein quantified”) and all missing values were turned to 0s (“protein not quantified”). Subsequently, treeClust dissimilarities were re-calculated based on this binary version of ProteomeHD. The performance of these different metrics was assessed by a precision - recall analysis as described above.

### Plasmids, siRNA, and antibodies

For cloning of peroxisome-targeted Miro1, the C-terminal TMD and tail of Myc-Miro1 (kindly provided by P. Aspenström, Karolinska Institute, Sweden) was exchanged by a PEX26/ALDP fragment previously shown to target proteins to the peroxisome membrane^82^. PEX11β-EGFP was kindly provided by G. Dodt (Univ. of Tuebingen, Germany). PEX11β siRNA (AUU AGG GUG AGA AUA GAC AGG AUGG) (Eurofins) was previously verified^110^. Control siRNA (si-GENOME nontargeting siRNA pool #2) was obtained from GE Healthcare (D-001206-14-05). Antibodies used were as follows: rabbit polyclonal antibody against PEX14 (1:1400, kindly provided by D. Crane, Griffith University, Australia); mouse monoclonal antibody 9E10 against the Myc epitope (1:200, Santa Cruz Biotechnology, Inc., sc-40), rabbit monoclonal antibody against PEX11β (1:1000, Abcam, ab181066); rabbit polyclonal antibody against GAPDH (1:2000, ProSci3783). Secondary anti-IgG antibodies against rabbit (Alexa 594, 1:1000, Molec. Probes/Life Technol. A21207) and mouse (Alexa 488, 1:400, Molec. Probes/Life Technol. A21202) were obtained from ThermoFisher Scientific. HRP-coupled donkey polyclonal antibody against rabbit IgG (1:5000) was obtained from Biorad (172-1013).

### Cell culture and transfection

COS-7 cells (African green monkey kidney cells; ATCC CRL-1651), and PEX5 deficient fibroblasts (kindly provided by H. Waterham, AMC, University of Amsterdam, NL) were cultured in DMEM (high glucose, 4.5 g/L) supplemented with 10% FBS, 100 U/ml penicillin and 100 μg/ml streptomycin at 37°C (5% CO_2_, 95% humidity) (HERACell 240i CO_2_ incubator). COS-7 cells were transfected using diethylaminoethyl-dextran (Sigma-Aldrich). dPEX5 fibroblasts have enlarged peroxisomes, which facilitates the visualization of membrane extensions. For transfection of dPEX5 fibroblasts, the Neon® Transfection System (Thermo Fisher Scientific) was used following the manufacturer’s protocol. Briefly, cells (seeded 24h before transfection) were washed once with PBS and trypsinized using TrypLE Express. Trypsinized cells were resuspended in complete medium, pelleted by centrifugation, and washed with PBS. The cells were once again centrifuged and carefully resuspended in 110 μl buffer R. For each condition, 4 × 10^5^ cells were mixed with the DNA construct (5 μg) or with 100 nM siRNA. Cells were microporated using a 100 μl Neon tip with the following settings: 1400 V, 20 ms, one pulse. Microporated cells were immediately seeded into plates with prewarmed complete medium (without antibiotics) and incubated at 37°C with 5% CO_2_ and 95% humidity. The efficiency of silencing was monitored by immunoblotting of cell lysates and confirmed as previously reported^110^.

### Immunofluorescence and microscopy

Cells grown on glass coverslips were processed for immunofluorescence 24h after transfection. Cells were fixed for 20 min with 4% paraformaldehyde in PBS (pH 7.4), permeabilized with 0.2% Triton X-100, and blocked with 1% BSA, each for 10 min. Incubation with primary and secondary antibodies took place for 1h each in a humid chamber. Coverslips were washed with ddH_2_O to remove PBS and mounted with Mowiol medium on glass slides. All immunofluorescence steps were performed at room temperature and cells were washed three times with PBS between each individual step. Cell imaging was performed using an IX81 microscope (Olympus) equipped with an UPlanSApo 100×/1.40 oil objective (Olympus). Digital images were taken with a CoolSNAP HQ2 CCD camera and adjusted for contrast and brightness using the Olympus Soft Imaging Viewer software and MetaMorph 7 (Molecular Devices). For live-cell imaging, COS-7 cells were plated in 3.5 cm diameter glass bottom dishes (Cellvis). MitoTracker Red CMXRos (Life Technologies) at 100 nM was used for visualisation of mitochondria. Live-cell imaging data was collected using an Olympus IX81 microscope equipped with a Yokogawa CSUX1 spinning disk head, CoolSNAP HQ2 CCD camera, 60 x/1.35 oil objective. Digital images were taken and processed using VisiView software (Visitron Systems, Germany). Prior to image acquisition, a controlled temperature chamber was set-up on the microscope stage at 37°C, as well as an objective warmer. During image acquisition, cells were kept at 37°C and in CO_2_ –independent medium (HEPES buffered). 200 stacks of 9 planes (0.5 µm thickness, 100 ms exposure) were taken in a continuous stream. All conditions and laser intensities were kept between experiments.

### Quantification and statistical analysis of peroxisome morphology and interaction

Analysis of statistical significance was performed using GraphPad Prism 5 software. A two-tailed unpaired *t* test was used to determine statistical difference against the indicated group. \**P* < 0.05, \*\**P* < 0.01, \*\*\**P* < 0.001. For analysis of peroxisome morphology, a minimum of 150 cells were examined per condition, and organelle parameters (e.g. membrane protrusions) were microscopically assessed in at least three independent experiments. The analysis was made blind and in different areas of the coverslip. Organelle interaction and contact time were analysed manually from live-cell imaging data using MetaMorph 7 (Molecular Devices). A region of interest (ROI) was drawn in different areas of the cell. Spherical and elongated peroxisomes within the ROI were tracked over the whole time course, and the frequency and duration of contacts monitored. Multiple interactions of the same peroxisome with mitochondria were treated as separate events. Data are presented as mean ± SD.

