## Supplementary Figures for "The human proteome co-regulation map reveals functional relationships between proteins"

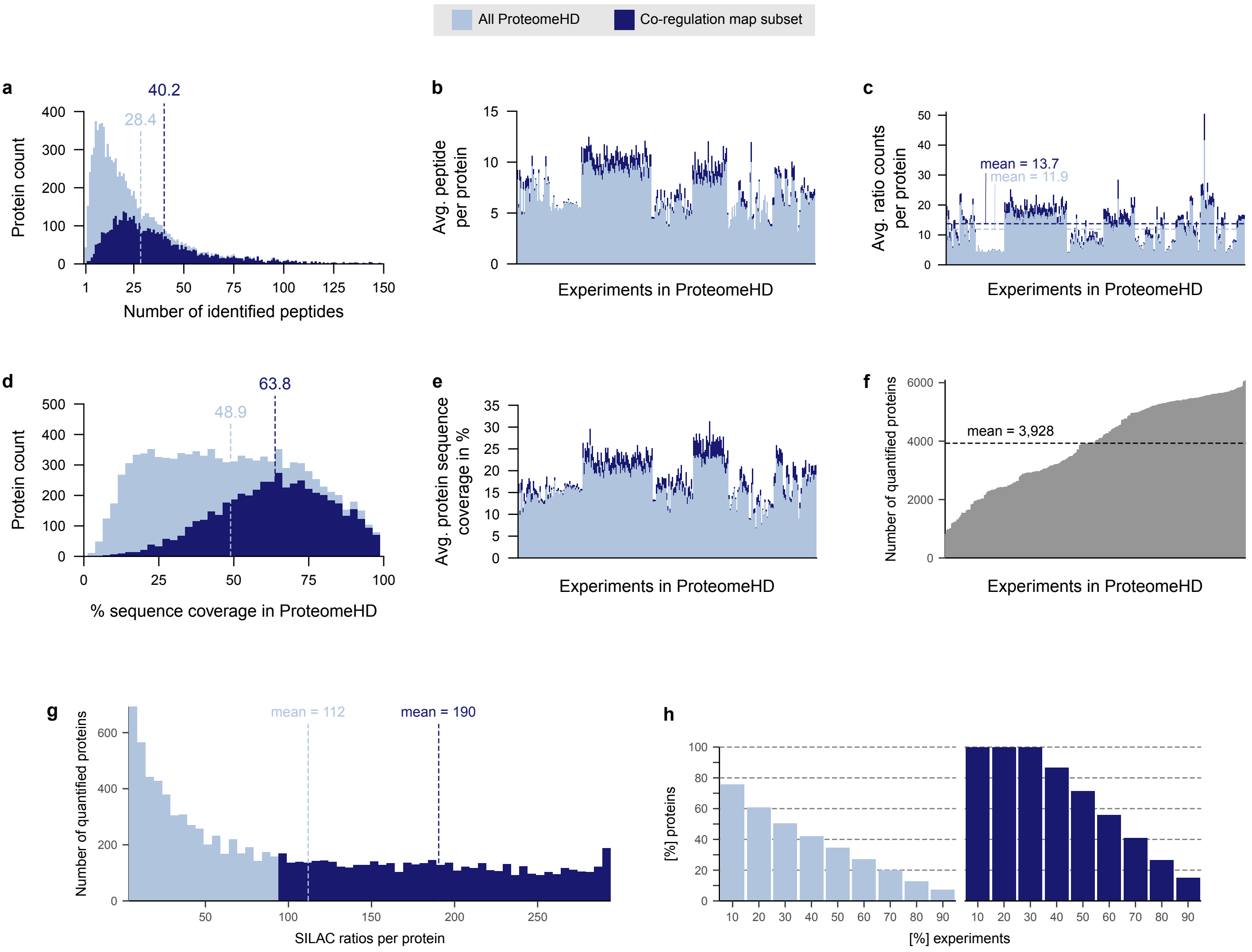


**Supplementary Figure S1. Peptide and protein quantitation statistics in ProteomeHD**

**(a)** Histogram showing the number of peptides identified per protein in ProteomeHD (10,323 proteins, light blue) and in the subset of ProteomeHD used to make the co-regulation map (5,013 proteins, dark blue). Dashed lines show the average number of peptides per protein. (**b**) Number of peptides per protein broken down by experiment. The average peptide number for the proteins detected in each experiment is shown. (**c**) Average number of SILAC ratio counts (independent observations) per protein, broken down into the 294 input experiments. (**d**) Sequence coverage of proteins in ProteomeHD. Dashed lines indicate the average. (**e**) Average sequence coverage of proteins in each input experiment. (**f**) The number of proteins that were quantified in the 294 experiments of ProteomeHD ranges from 817 to 6,080. The average is 3,928 proteins per SILAC ratio. (**g**) Number of experiments, i.e. SILAC ratios, in which proteins were quantified. Only proteins that were quantified in at least 95 experiments were used for the co-regulation analysis. On average, proteins in ProteomeHD were quantified in 112 input experiments. The average rises to 190 if only proteins used for the co-regulation analysis are considered. (**h**) Barchart showing which fraction of proteins have been detected in which fraction of experiments. For example, 100% of proteins in the co-regulation map have been quantified in at least 30% of the 294 experiments. About 15% of the proteins have been quantified in at least 90% of the experiments.

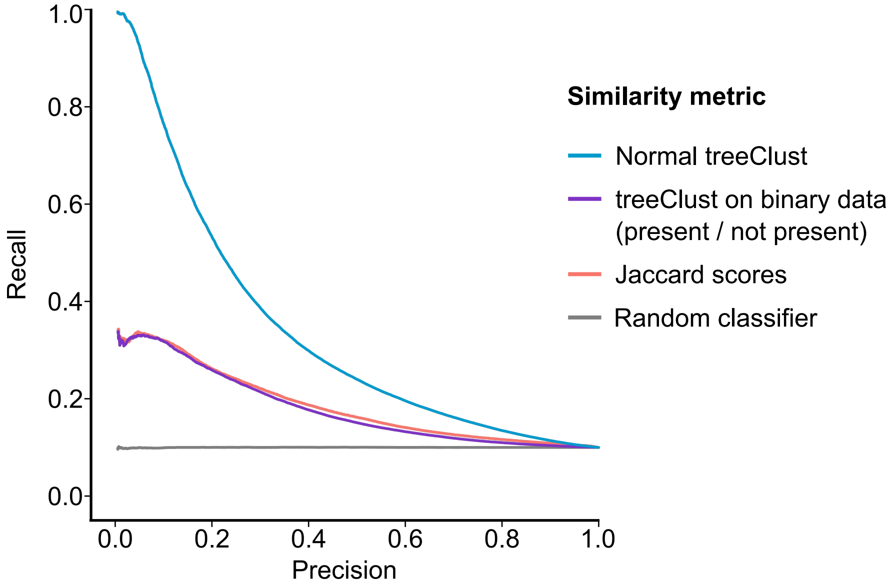


**Supplementary Figure S2. Impact of co-occurrence on treeClust learning**

Performance comparison of a standard treeClust application on ProteomeHD with two types of co-occurrence measures. Jaccard scores are an established co-occurrence measure (protein pairs observed in the same set of experiments would get a Jaccard score of 1, while protein pairs without any overlapping experiments would get a score of 0). We also applied treeClust to a "binary" version of ProteomeHD, where all SILAC ratios were set to 1 and all missing values were set to 0. The precision recall curve uses Reactome as a gold standard. It shows that Jaccard and "binary treeClust" work equally well but both are outperformed by the standard co-regulation analysis. Therefore, while co-occurrence of proteins across ProteomeHD does provide some information about functional associations, quantitative up- and downregulation is a far better indicator of shared protein function, at least for ProteomeHD. Notably, this also shows that treeClust can detect co-occurrence in principle, if the data are transformed into a binary format.

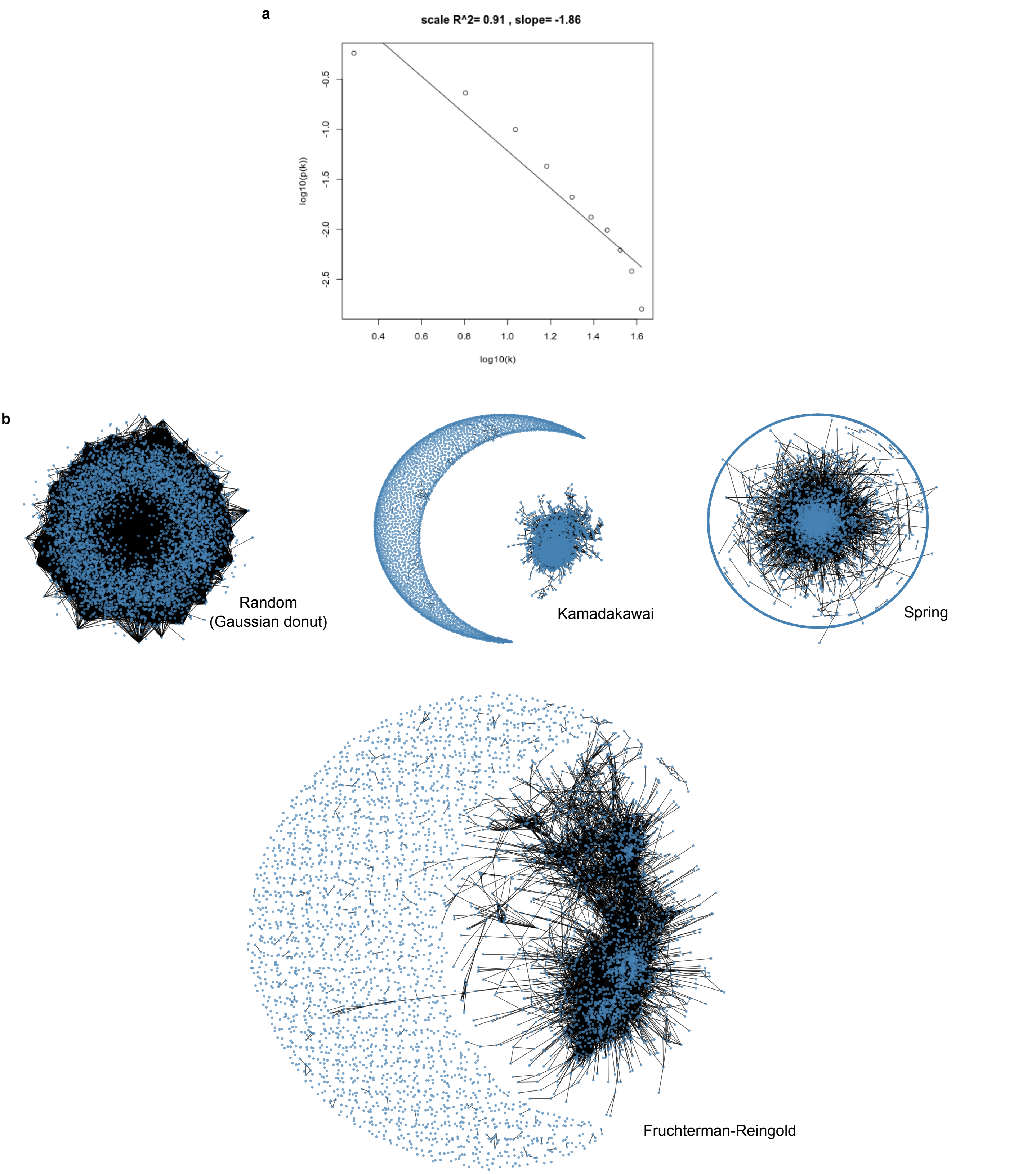


**Supplementary Figure S3. The protein co-regulation network satisfies scale-free topology but is difficult to visualize as an interaction network**

(**a**) The "scale free plot" produced by the WGCNA R package using the treeClust-derived adjacency matrix. The log of the connectivity k is plotted against the log of the frequency of this connectivity. There is a linear relationship between these two variables, as indicated by the square of the Pearson correlation, R2, being 0.91. This shows that the protein co-regulation network derived from ProteomeHD using treeClust is at least approximately scale free. (**b**) Visualization of a weighted, undirected network with 5,013 nodes (proteins detected in at least 95 experiments) and 62,812 edges (top scoring 0.5% of links), based on the co-regulation score. Four common algorithms were used to create different network layouts, but with so many edges it is difficult to avoid the "hairball" problem.

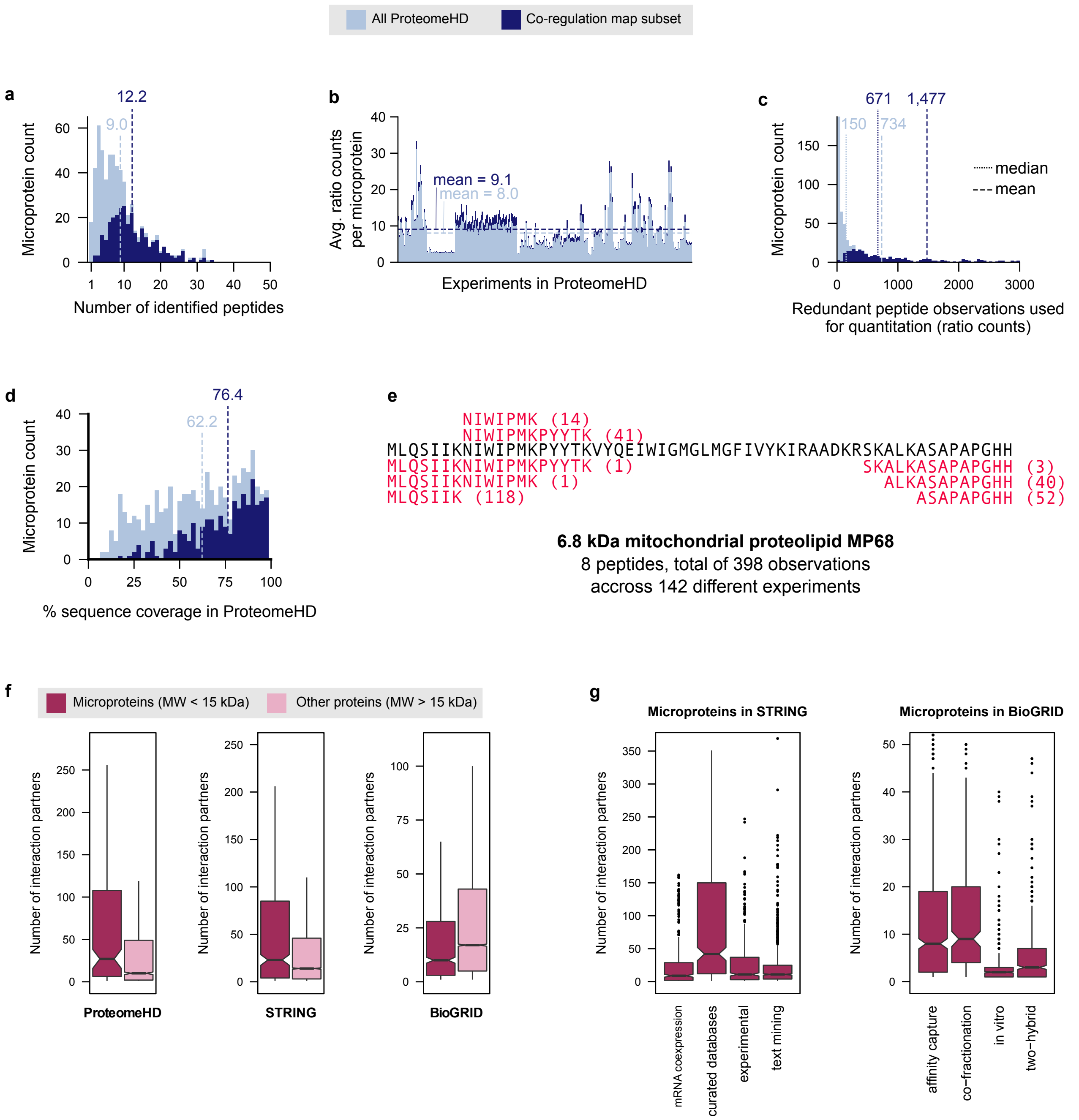


**Supplementary Figure S4. Microproteins in ProteomeHD and their connectivity**
(**a**) Histogram showing the number of peptides identified per microprotein (proteins < 15 kDa) in ProteomeHD and the subset of ProteomeHD used to make the co-regulation map. Dashed lines show the average number of peptides per microprotein. (**b**) Average number of SILAC ratio counts (independent observations) per microprotein, broken down into the 294 input experiments. (**c**) Histogram showing the cumulative SILAC ratio counts per microprotein across all experiments in ProteomeHD. (**d**) Sequence coverage of microproteins in ProteomeHD. Dashed lines indicate the average. (**e**) The actual peptides for one example microprotein, MP68. The numbers in brackets indicate in how many different experiments each peptide was observed. **(f)** Microproteins tend to have more co-regulation partners in ProteomeHD than larger proteins (median 27 *vs* 10 associations). Microproteins also have more functional protein - protein associations according to STRING (median 23 *vs* 14). However, larger proteins have considerably more physical interaction partners than microproteins, according to BioGRID (median 10 *vs* 17). (**g**) The number of interaction partners of microproteins identified by STRING and BioGRID, broken down by the evidence type available in each resource. We considered STRING interactions with a minimum score of 400 in the individual evidence channels (e.g. mRNA coexpression). Two STRING evidence channels (gene neighborhood and evolutionary co-occurrence) were omitted because they contribute very little. For panel (f) we considered only the most reliable STRING interactions, i.e. those with a combined interaction score above 900.

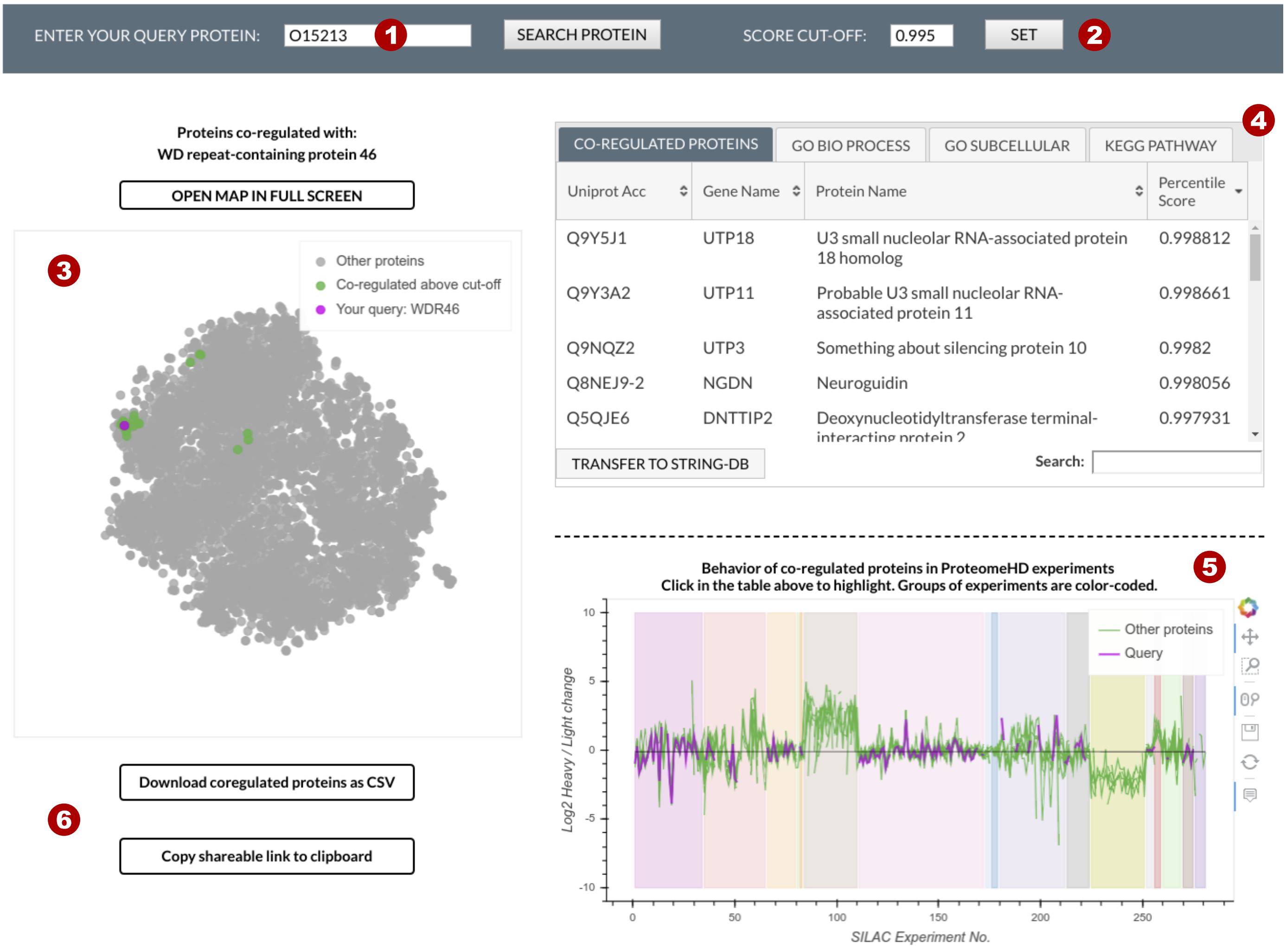


1 **Search field.** Search for a protein of interest using Uniprot IDs or gene names. Suggestions of proteins that are actually in the map appear upon typing.

2 **Set co-regulation score cut-off.** Here it is expressed as a percentile score to allow for a more intuitive exploration. For example, a cut-off of 0.9 shows associations that are stronger then 90% of all associations detected in the co-regulation analysis of ProteomeHD. Defaults to 0.995 (showing the top-scoring 0.5% pairs, as in the manuscript).

3 **Co-regulation map.** Zoom using mouse wheel. Protein names appear when hovering mouse over points. Query protein highlighted in purple, proteins co-regulated with it - above the chosen cut-off - are shown in green. These may not be adjacent to query protein in the map, as they may be co-regulated with other proteins as well.

4 **Functional characterisation of co-regulation partners.** Shows list of proteins co-regulated with query protein above the selected cut-off. Bottom left button sends the list to STRING. Separate tabs show enrichment of GO terms and KEGG pathways among the co-regulated proteins, together with Bonferroni - adjusted p-values.

5 **Co-regulation patterns.** Line plots showing up- and down-regulation of the query protein and up to 100 co-regulated proteins across ProteomeHD. Zoom in using mouse wheel. Background colours distinguish different input projects, project names are shown when hovering mouse over them.

6 **Save the results.** To save or share the results, one can either download a csv file or copy a link that will reproduce the current selection of query protein and score cut-off.

**Supplementary Figure S5. Layout of www.proteomeHD.net**
Screenshot of the core page of www.proteomeHD.net, an interactive web-based app to explore co-regulation data. The basic elements are highlighted and explained. Note that the page also contains help and download sections.

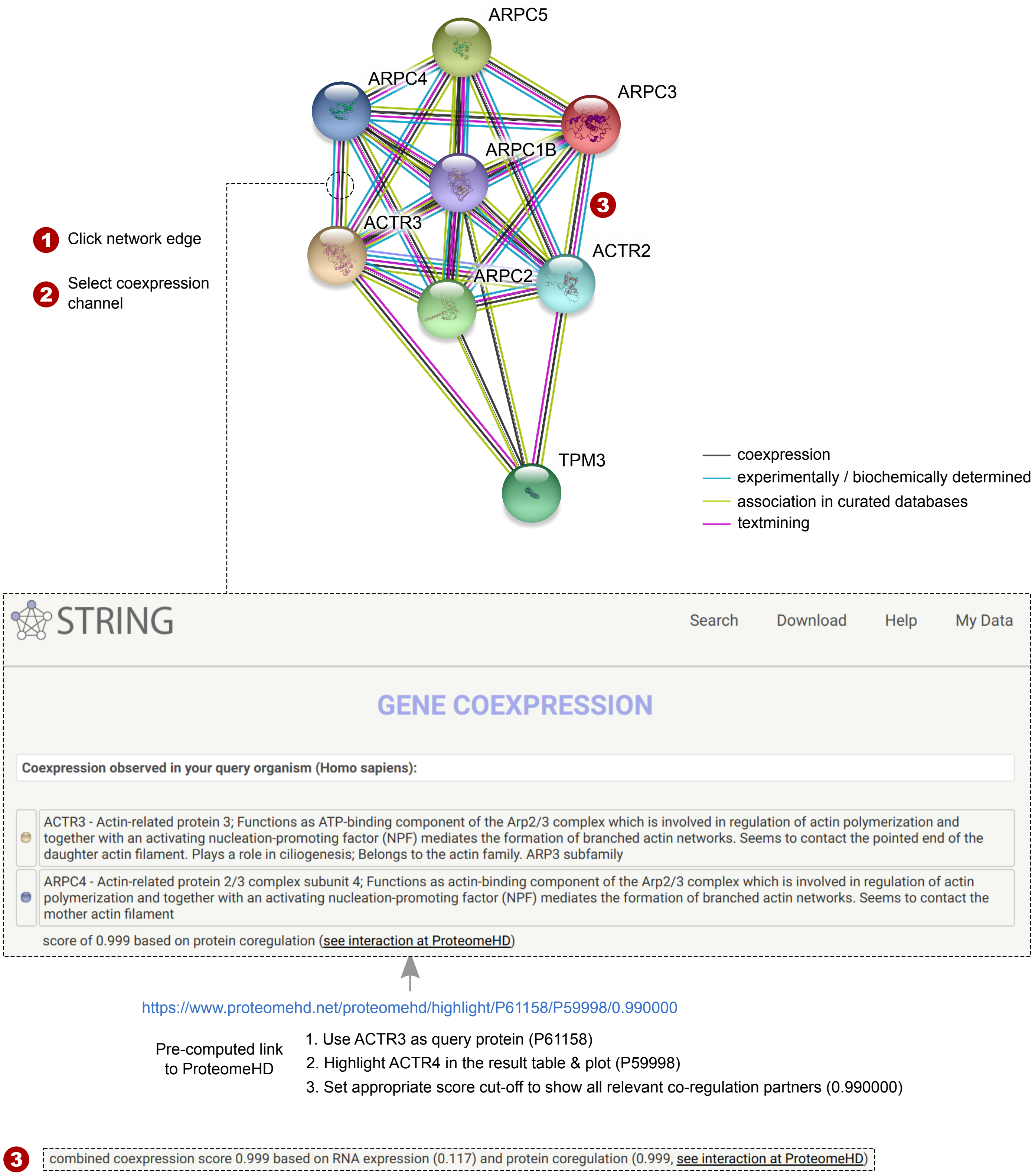


**Supplementary Figure S6. Integration of co-regulation scores with STRING (https://string-db.org)**
A typical protein - protein association network in STRING, containing the Arp2/3 complex and tropomyosin 3, both of which are involved in actin cytoskeleton regulation. Network edges are colour-coded by the type of evidence available for the association. Protein co-regulation information is embedded in the gene coexpression channel. The channel view shows the channel-specific STRING score, a re-calibrated version of our co-regulation score. It also contains a pre-computed link to www.proteomeHD.net, which uses the first protein as ProteomeHD query and highlights the second protein in the results. If more than one protein isoform is available in ProteomeHD, STRING will link to the alphabetically first isoform, which is generally the main one. The link also contains a cut-off setting to match the ProteomeHD cut-off to the equivalent one selected by the user in STRING. In cases where both mRNA coexpression and protein co-regulation evidence is available for an association, their relative contribution to the STRING coexpression score is indicated (shown here as point 3).

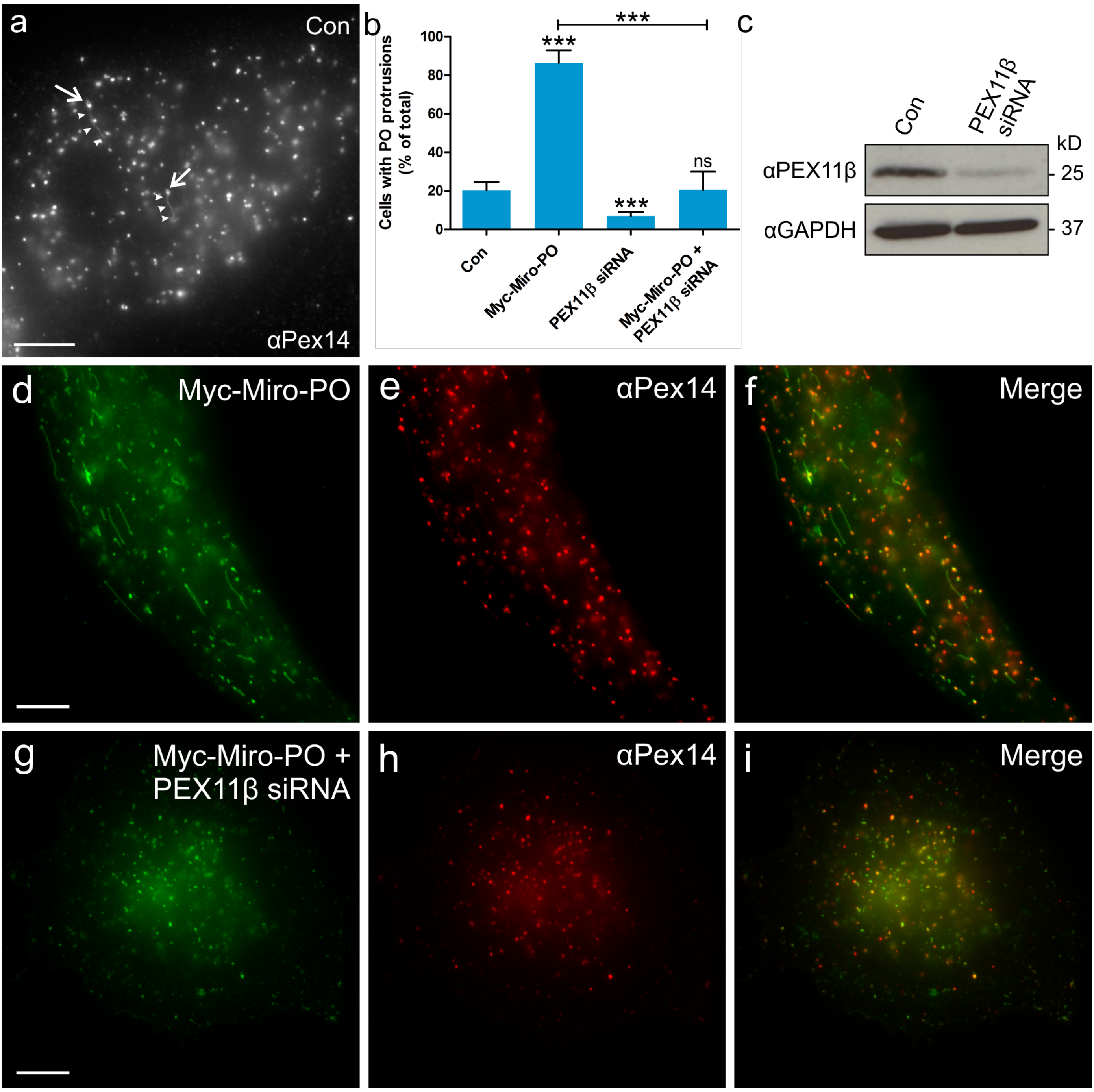


**Supplementary Figure 7. MIRO1-induced peroxisomal membrane protrusions depend on PEX11β**

(**a-i**) PEX5-deficient human skin fibroblasts were mock-treated (control), or transfected with Myc-Miro-PO, a peroxisome-targeted Miro1 variant, in the presence of control- or PEX11β-specific siRNA. Cells were processed for immunofluorescence using anti-Myc and anti-PEX14 antibodies (peroxisomal marker). (**b**) Quantification of cells with peroxisomal protrusions. The average result of 3 independent experiments is shown, error bars indicate standard deviation. (**c**) Immunoblots of cell lysates showing efficient silencing of Pex11β. Loading control: GAPDH. (**a, b**) Control cells occasionally contain peroxisomes with membrane protrusions (< 5 per cell; up to 5 µm in length). (**d-f**, **b**) Myc-Miro-PO induces the formation of peroxisomal membrane protrusions (> 5 per cell; > 5 µm in length). (**g-i**, **b**) Silencing of PEX11β by siRNA significantly reduces the number of cells with peroxisomal membrane protrusions in controls and Myc-Miro-PO expressing cells. Globular peroxisomes (arrows) with membrane protrusions (arrowheads) in (a) are highlighted. *** $P < 0.001$; ** $P < 0.01$ from a two-tailed unpaired t test; ns, not significant. Scale bars, 10 µm.

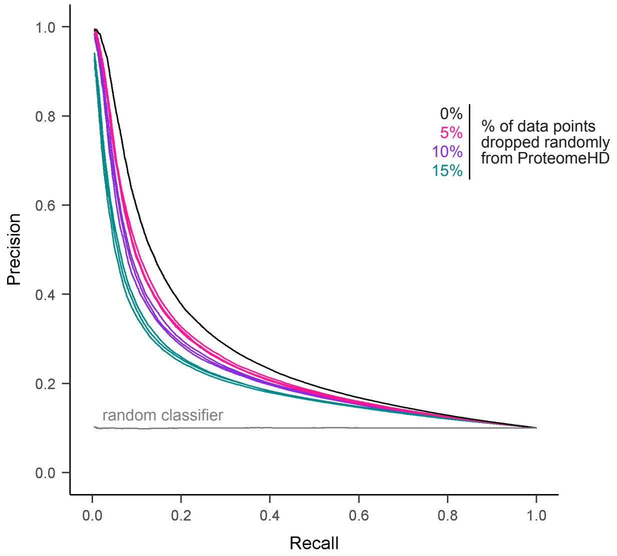


**Supplementary Figure 8. Information content of ProteomeHD has not reached saturation yet**
We randomly removed 5%, 10% and 15% of the data points across the ProteomeHD matrix, in triplicate, and repeated treeClust learning to predict protein associations. The Precision-Recall analysis shows that removing data points decreases performance proportionally to the amount of removed data, suggesting that adding additional data would likely enhance performance further.

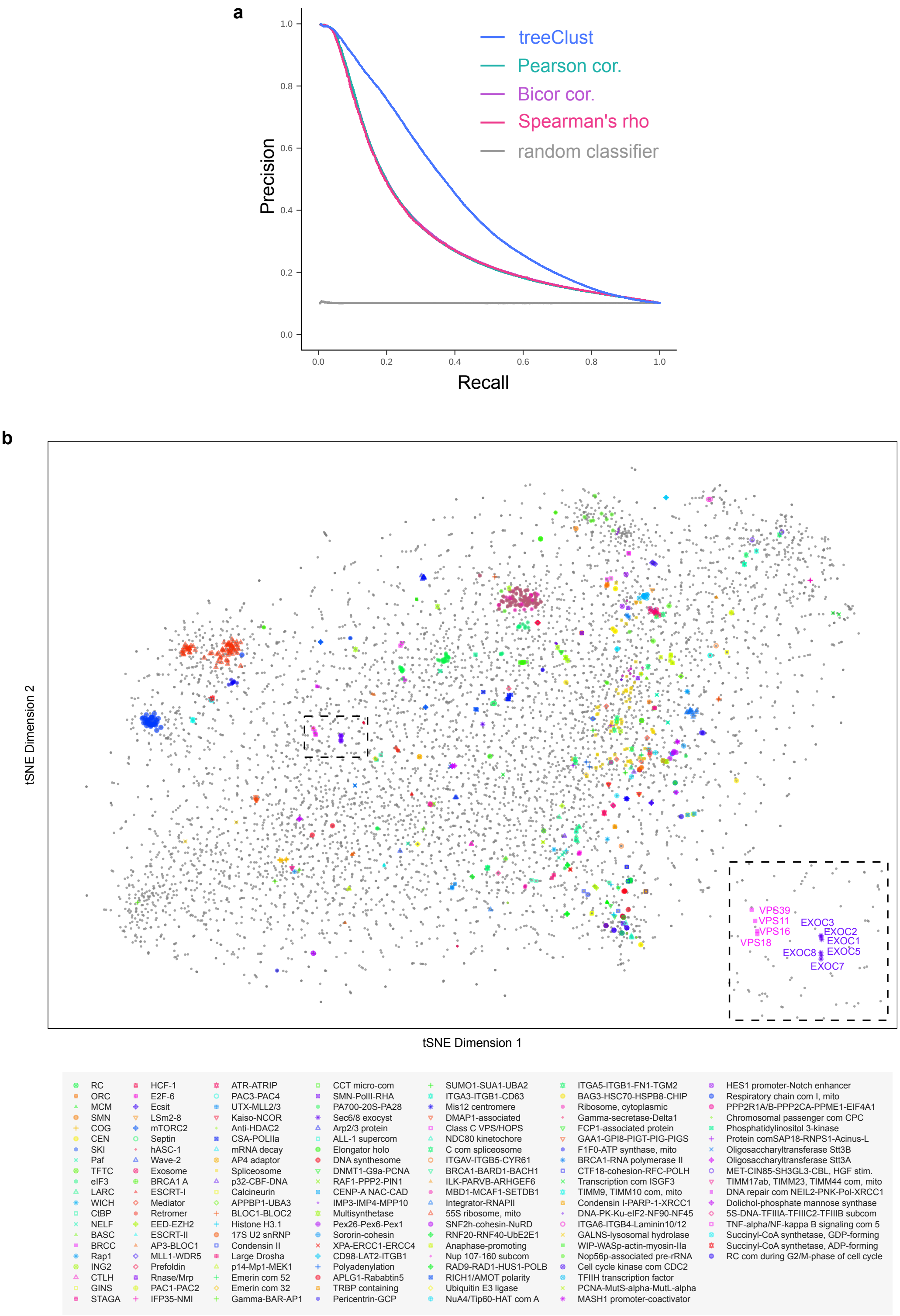


**Supplementary Figure 9. Validation of treeClust and t-SNE on an independent proteomics dataset**
(**a**) treeClust was applied to the TMT-based cancer proteomics dataset from Lapek *et al* (ref. 20). It outperforms Pearson, Spearman and Bicor correlation, as shown by a Precision-Recall analysis using Reactome annotations as the gold standard. Note that treeClust builds only one decision tree per condition, i.e. 41 trees on this dataset, too few for a standard analysis. Therefore, treeClust was performed iteratively, obtaining the mean co-regulation score of 100 treeClust forests, each generated from 10 random experiments. (**b**) Co-regulation map for the Lapek *et al* dataset, made by t-SNE from treeClust scores. As in the correlation network of the original report (Fig. 2 in ref 20), CORUM protein complexes are colored. In contrast to a network, there is not a limited number of arbitrarily arranged, pairwise links, but the position of each protein reflects its similarity or dissimilarity to all other proteins in the map. This makes it possible to place all proteins in a functional context, not just those that are directly linked to members of the core network. It also allows for a hierarchical analysis of protein associations, with increasing distances indicating weaker co-regulation. For example, the subunits of the protein complexes in the enlarged map area (inset) are clustered together, and the distances between the complexes are larger. However, all complexes have roles in vesicular trafficking.
